# SET1B drives sustained HIF activity and disease progression in clear cell renal cell carcinoma

**DOI:** 10.1101/2025.05.13.653699

**Authors:** Brian M Ortmann, James Bertlin, Rachel Seear, Esther Arnaiz, Salwa Lin, Ahmed M Salman, Laura Wilson, Adrian L Harris, Grant D Stewart, Anatasia Hepburn, Simon Clare, Craig N Robson, Anneliese O Speak, James A Nathan

## Abstract

The cellular response to hypoxia is driven by hypoxia-inducible factors (HIFs), which regulate genes involved in glycolysis, angiogenesis, and cell proliferation, as well as inflammation and tumour progression. HIF activation is well-characterised and is primarily regulated by oxygen-dependent prolyl hydroxylation and subsequent degradation. However, how transcription of individual HIF target genes is regulated at the chromatin level is less clear. SET1B, a histone H3 lysine 4 (H3K4) methyltransferase, has emerged as a key modulator of HIF target gene transcription. Our study reveals that SET1B interacts with RNA Polymerase II to coordinate sustained HIF-mediated transcriptional activity through multiple functional domains. We also show that in clear cell renal cell carcinoma (ccRCC), SET1B is critical for sustained HIF activity, and SET1B expression correlates with disease progression and metastasis in patient samples. Moreover, SET1B depletion enhances the efficacy of HIF-2 inhibitors, establishing SET1B as a potential therapeutic target in ccRCC.

## Introduction

Cellular responses to hypoxia are essential for the survival of multicellular organisms and involve transcriptional reprogramming driven by the hypoxia-inducible factor (HIF) family of transcription factors^1^. HIFs regulate genes critical for glycolysis, angiogenesis, and cell proliferation^2^. Moreover, HIFs also have roles in inflammation, immune modulation and tumour progression, making the HIF pathway a promising therapeutic target^3–5^.

HIF regulation primarily occurs post-transcriptionally via oxygen-dependent degradation of the two main HIF-α isoforms, HIF-1α and HIF-2α. Under normoxic conditions, HIF undergoes oxygen-dependent prolyl hydroxylation by prolyl hydroxylase domain (PHD) enzymes, facilitating the binding of the von Hippel-Lindau (VHL) E3 ubiquitin ligase complex, leading to ubiquitination and proteasomal degradation^6–10^. In hypoxic conditions, decreased PHD activity stabilises HIF-α, promoting dimerisation with HIF1β to form a heterodimer complex that activates gene transcription.

While the mechanisms underlying HIF activation are well characterised, how HIFs transcriptionally regulate individual target genes across diverse cellular and pathological contexts remains an important area of investigation. This has significant clinical implications, as targeting HIF presents an attractive therapeutic strategy.

However, the potential toxicity associated with broadly suppressing HIF activity remains a challenge. Transcriptional co-activators and epigenetic modifications have been shown to facilitate HIF specificity, offering an alternative approach to target specific pathways of HIF activity^11–13^. Early studies identified CBP/p300 as co-activators for subsets of HIF target genes^14^, while more recent research has uncovered roles for other co-activators, including TIP60, CDK8 Mediator, DNA-PK, and the FACT complex^15–18^. HIF must also interact with the transcriptional machinery, as transcriptional outcomes are largely dependent on RNA Polymerase II (Pol II) activity^19,20^. An accepted role of transcriptional activators is to promote Pol II recruitment and initiation. However, recent evidence suggests that Pol II often initiates transcription but pauses shortly downstream of the transcriptional start site, a phenomenon observed in 40-70% of genes involved in stress responses, including hypoxia^21–23^. Therefore, how HIF and associated transcriptional activators interact with Pol II is not well understood.

We recently discovered that the SET1B histone 3 lysine 4 (H3K4) methyltransferase plays a crucial role in regulating the transcription of HIF target genes, particularly those typically expressed at low levels under normoxic conditions^24^. While this effect of SET1B could be attributed to increased histone methylation during hypoxia, given that H3K4 methylation is traditionally associated with active transcription^25^, H3K4me3 levels did not consistently correlate with reduced mRNA expression. In fact, depletion of SET1B led to a reduction in H3K4me3 levels across gene bodies rather than being limited to transcriptional start sites of specific HIF target genes, indicating that SET1B plays a broader role in modulating HIF transcriptional activity. SET1B also contains additional functional domains beyond its known SET enzymatic domain, but how these regions contribute to HIF-mediated transcription is not known.

Here, we demonstrate that multiple functional domains of SET1B, including its interaction with RNA Pol II, are critical for modulating HIF transcriptional activity under hypoxic conditions. Using a model of sustained HIF activation, we also show that SET1B is essential not only for initiating HIF activity but for maintaining its prolonged activation. These findings have direct relevance to human disease, as using clear cell renal cell carcinoma (ccRCC) as cancer type where sustained HIF activity is important for tumorigenesis, we find that SET1B is indispensable for maintaining HIF activity, with its expression correlating strongly with disease progression in patient samples. Furthermore, SET1B depletion enhances the efficacy of a clinically relevant HIF-2 inhibitor and reduces HIF-dependent transcription, even in cases of resistance to HIF-2 inhibitors. These findings establish SET1B as a driver of HIF-dependent ccRCC progression and emphasise its potential as a therapeutic target in ccRCC.

## Results

### SET1B interacts with the RNA Pol II complex and requires multiple functional domains to regulate HIF activity

Our previous studies identified SET1B as a selective mediator of HIF target gene expression in hypoxia^24^ (**Figure S1A**). However, while loss of SET1B reduced H3K4me3 levels at specific HIF targets, transcriptional changes were not always proportional to the reduction in H3K4me3, suggesting additional roles for SET1B beyond its methyltransferase activity. We therefore set out to determine whether other SET1B functional domains contribute to HIF transcriptional regulation.

SET1B is a member of the evolutionarily conserved family of H3K4 methyltransferases that includes MLL1-4 and SET1A, (also referred to as the COMPASS family)^26–28^. Each methyltransferase forms large complexes that share core subunits ASH2L, RBBP5, WDR5, and DPY30 to modify histone tails.

Distinctively, SET1A and B contain additional regulatory subunits, CFP1 (CXXC1) and WDR82, which aid in promoter recruitment and facilitate interactions with the RNA Pol II complex, respectively^29,30^. SET1B also comprises internal functional domains: the catalytic SET domain for H3K4me3 deposition, an uncharacterised RNA-recognition motif (RRM), and a DPR motif that mediates RNA Pol II interaction via WDR82^31,32^ (**Figure 1A**). We therefore generated mutants deficient in these SET1B functional regions (ΔRRM, ΔSET and a DPR to AAA mutant), and expressed these in HEK293T cells to examine whether: (1) whether a SET1 complex can still form, and (2) determine if these SET1 mutants alter HIF transcriptional signalling.

**Figure 1.**
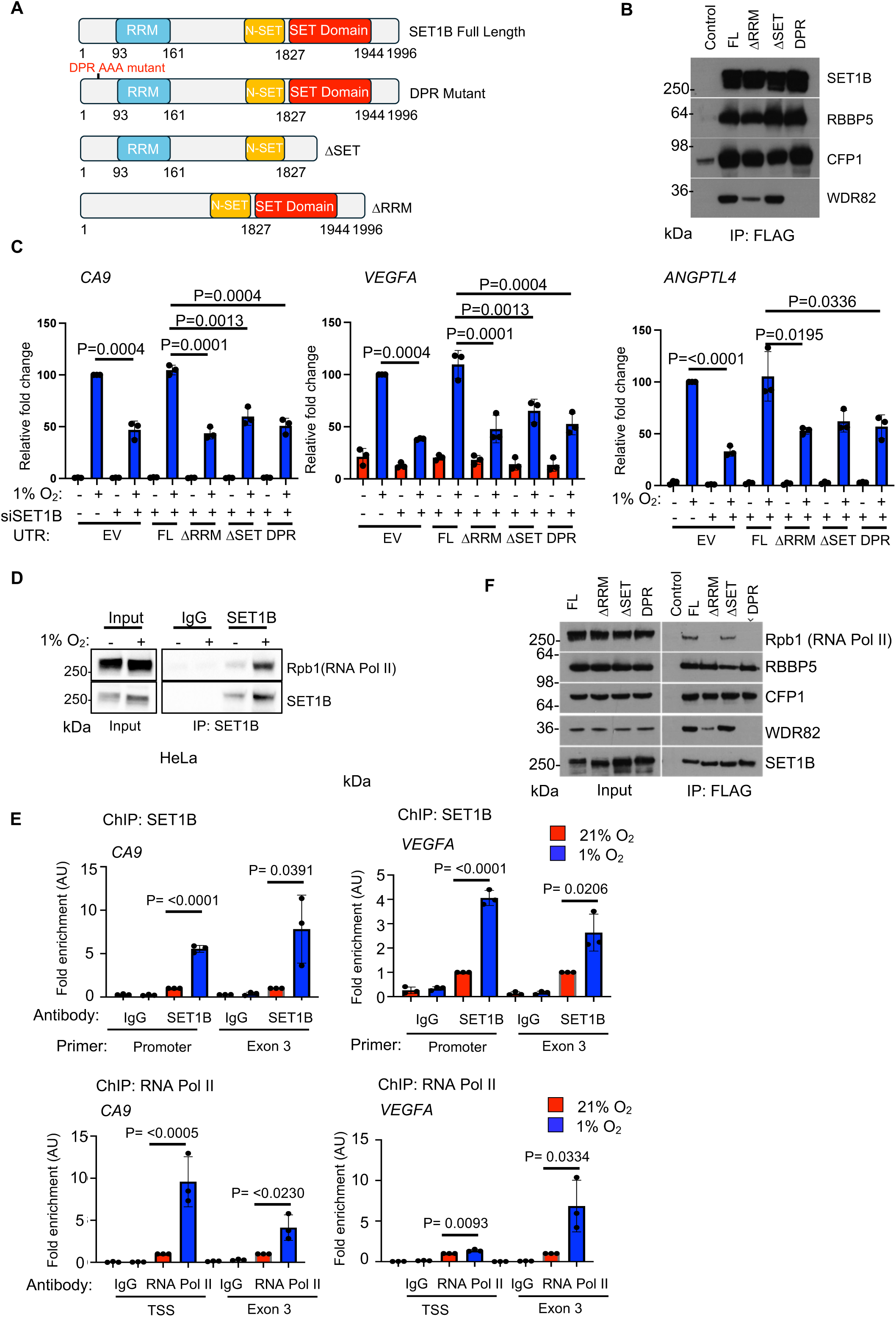
SET1B interacts with the RNA Pol II complex and requires multiple functional domains to coordinate HIF activity. **(A)** Schematic representation of full-length SET1B and truncation mutants. Key functional domains are highlighted, including the SET domain (associated with methylation activity), RNA recognition motif (RRM), and DPR motif (involved in RNA Pol II binding). **(B)** HEK293T cells were transfected with 4 µg of the indicated expression constructs for 48 hours. Constructs were immunoprecipitated using FLAG-magnetic beads, and their interaction with other members of the SET complex was evaluated by immunoblotting. **(C)** A549 cells stably expressing full-length SET1B or SET1B truncation mutants were transfected with a SET1B siRNA targeting the 3′UTR and incubated under 21% or 1% oxygen for 24 hours. mRNA expression of HIF target genes (*CA9*, *VEGFA*, *ANGPTL4*) was assessed using qPCR. Graphs represent the mean ± SD from three biological replicates. Statistical analysis was performed using two-way ANOVA. **(D)** Co-immunoprecipitation of SET1B and RNA Pol II. SET1B was immunoprecipitated from HeLa cells incubated in 21% or 1% oxygen for 6 hours. Samples were analysed by immunoblotting with the indicated antibodies. Results are representative of three biological replicates. **(E)** HEK293T cells were transfected with 4 µg of the indicated expression constructs for 48 hours. Constructs were immunoprecipitated using FLAG-magnetic beads, and their interaction with other members of the SET complex and RNA Pol II was assessed by immunoblotting. **(F)** ChIP-PCR analysis of SET1B and RNA Pol II in A549 cells exposed to 21% or 1% oxygen for 6 hours. Primers targeted the promoter and exon 3 regions of the *CA9* and *VEGFA* genes. Data represent the mean ± SD from three biological replicates. Statistical significance was determined using two-way ANOVA.

Immunoprecipitation experiments revealed that the ΔSET mutation did not impair SET1B’s ability to form a functional complex. In contrast, the ΔRRM reduced interactions between SET1B and WDR82, while the DPR mutant abolished binding to WDR82 (**Figure 1B**), consistent with prior findings^32^. Notably, interactions with other core subunits, such as CFP1 and RBBP5, remained intact in these mutants, demonstrating that loss of these domains did not affect SET1B’s ability to form a functional complex.

To investigate the functional significance of these SET1B mutants in regulating HIF activity, we depleted endogenous SET1B (using an siRNA targeting its 3′ untranslated region (UTR)) and stably expressing wild-type or mutant SET1B in A549 lung adenocarcinoma cells (**Figure 1C; S1B-E**). Depletion of endogenous SET1B reduced hypoxic induction of the HIF target genes, (*CA9, VEGF* and *ANGPTL4*), and the siRNA targeting the SET1B 3’ UTR reduced SET1B levels comparable to a siRNA pool targeting the SET1B coding sequence (**Figure S1C and D**). Reconstitution with wild-type SET1B restored expression of the HIF target genes under hypoxia, consistent with its requirement for facilitating HIF target gene expression, whereas all the SET1B mutants failed to rescue HIF transcriptional activity (**Figure 1C**).

We next explored whether the association between RNA Pol II and SET1B was altered by these different SET1B mutations, given that the DPR motif mediates SET1B’s interaction with RNA Pol II^32^. Endogenous SET1B interacted with the RNA Pol II complex subunit Rbp1 under both normoxia (21% oxygen) and hypoxia (1% oxygen) in HeLa cells (**Figure 1D**), but interestingly, this interaction was enhanced under hypoxic conditions, potentially reflecting increased chromatin recruitment of SET1B from the cytoplasm in hypoxia^24,33^. Chromatin immunoprecipitation (ChIP) assays revealed that SET1B was not only present at gene promoters, but that both SET1B and RNA Pol II bound the gene bodies of HIF target genes (*CA9* and *VEGFA*) under hypoxia (**Figure 1E**), consistent with SET1B associating with RNA Pol II during transcriptional elongation. The interaction between RNA Pol II and SET1B was dependent on the DPR motif, as expected^32^, with the DPR mutant completely disrupting SET1B’s association with both RNA Pol II and WDR82 (**Figure 1F**). Loss of the RRM also reduced interaction with RNA Pol II (**Figure 1F**), suggesting that the RRM domain plays a critical role in stabilising the interaction between RNA Pol II and SET1B through WDR82. However, the ΔSET mutation did not disrupt RNA Pol II binding (**Figure 1F**), but was still required to restore HIF-dependent transcription (**Figure 1C**). Therefore, these findings indicate that both the SET region and the ability of SET1B to bind to RNA Pol II are important for the regulation of HIF-mediated transcription.

### SET1B is required for sustained HIF-2-dependent activity in ccRCC

As SET1B binding was detected within the gene body of HIF target genes, we hypothesised that SET1B may be equally important for sustaining HIF activity after HIF binding and initiation of the hypoxic response. To explore this, we utilised clear cell renal cell carcinoma (ccRCC) as a model of sustained HIF activation caused by the loss of VHL function^34,35^. Of note, ccRCC typically demonstrates sustained HIF-2 activation, as HIF-1 is frequently deleted as an early tumour initiating event^36^.

We depleted SET1B in a panel of ccRCC cell lines (786-0, 769-P, RCC4, and RCC10) using siRNA, and measured the expression of HIF target genes (**Figure 2A**). SET1B depletion in all cell lines led to a selective reduction in HIF target gene expression, with effects on *VEGFA* and *ANGPTL4* but not *GLUT1*, as expected^24^. Similar results were obtained using shRNA- and CRISPR-mediated depletion of SET1B in 786-0 cells (**Figure S2A, B**). Although SET1B depletion reduced HIF dependent transcription across all cell lines, the effect was less pronounced in 769-P cells. Such differences in sensitivity are to be expected and are consistent with differential responses of ccRCC cells to HIF-2α inhibition^37^. We also confirmed that the effect of SET1B depletion was HIF-dependent, as HIF-2α knockout cells showed no further reduction in HIF target gene transcription in the absence of SET1B (**Figure S2C**). Immunoprecipitation experiments confirmed that endogenous SET1B interacts with HIF-2α in 786-0 cells (**Figure 2B**). Furthermore, we mapped the interaction between HIF-2α and SET1B, revealing that SET1B preferentially binds to the C-terminus of HIF-2α, specifically interacting with the PAS-B domain and the C-terminus, with no detectable interaction with the N-terminus, which includes the basic helix-loop-helix (bHLH) domain and PAS-A domain (**Figure S2D**).

**Figure 2.**
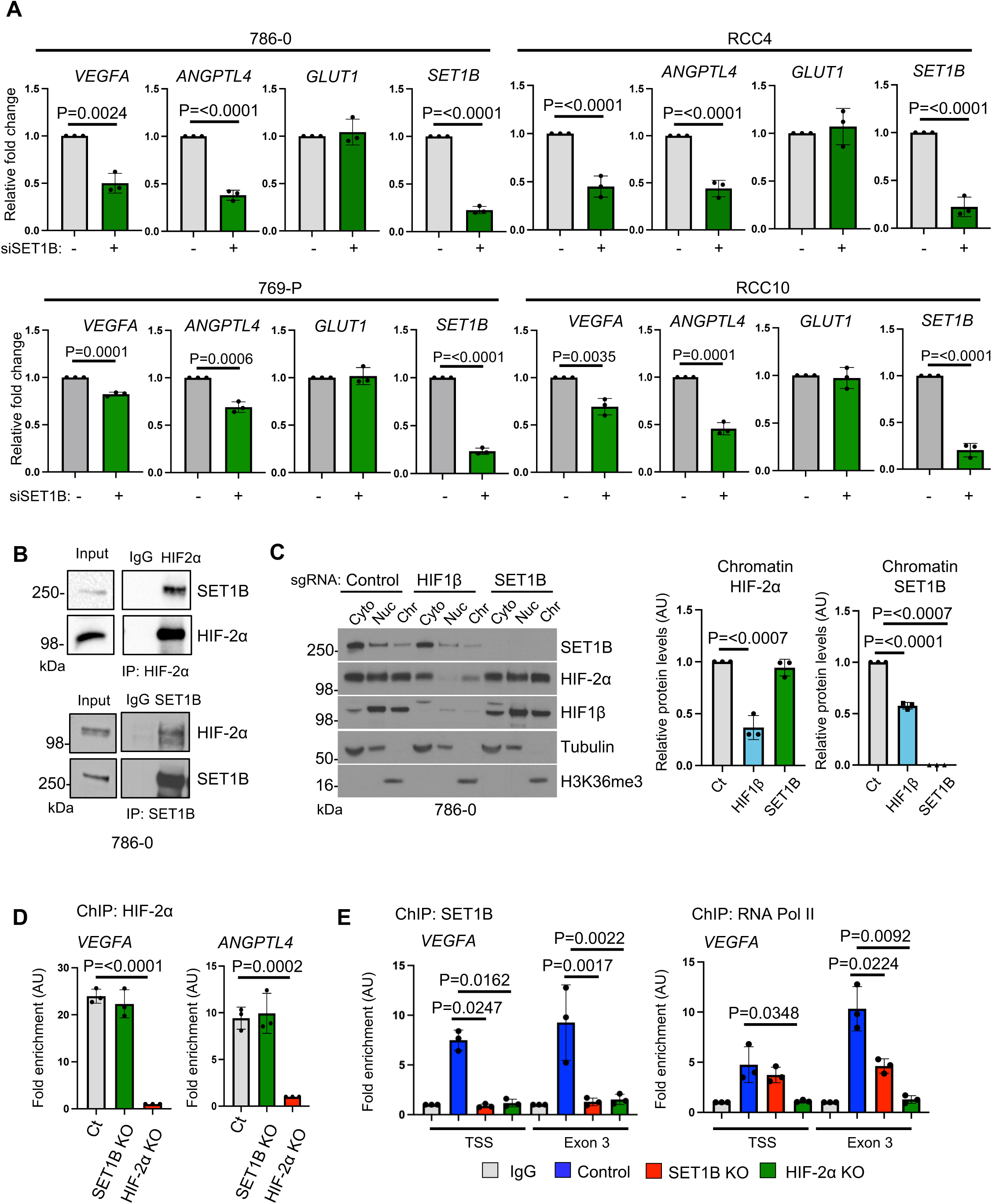
SET1B is required for sustained HIF activity. **(A)** 786-0 and RCC4, 769-P and RCC10 ccRCC cell lines were transfected with control or SET1B siRNA for 48 hrs. The expression of HIF target genes and *SET1B* was assessed using qPCR. Graphs are representative for 3 biological replicates and depict the mean ± SD. Two-way ANOVA. **(B)** Endogenous HIF-2α or SET1B was immunoprecipitated from 786-0 cells and immunoblotted. Immunoblot is representative of 3 biological replicates. **(C)** 786-0 cells were fractionated into cytoplasmic, nucleoplasm and chromatin fraction and immunoblotted for SET1B and HIF2-α. The ratio of SET1B and HIF-2α in the chromatin fraction was quantified using ImageJ (*n* = 3 biologically independent experiments, mean ± SD). **(D)** ChIP- PCR of HIF-2α in wildtype, SET1B KO and HIF-2α KO 786-0 cells. ChIP-PCR was performed using primers targeting the promoters of *VEGFA* and *ANGPTL4.* Graphs are representative for 3 biological replicates and depict the mean ± SD. Two-way ANOVA. **(E)** ChIP-PCR of SET1B and RNA Pol II in wildtype, SET1B and HIF-2α KO 786-0 cells. ChIP-PCR was performed using primers targeting the promoter and Exon 2 of *VEGFA* (n= 3 biological replicates). Graphs show mean ± SD. Two-way ANOVA.

Our previous work demonstrated that under hypoxic conditions, SET1B re-localises from the cytosol to chromatin in a HIF-dependent manner^24^. However, whether sustained HIF activation alters the subcellular localisation of HIF-2α and SET1B had not been determined. To address this, we performed subcellular fractionation in wild-type, HIF1β, or SET1B knockout (KO) 786-0 cells. We found that SET1B was distributed between the cytoplasm, nucleus, and chromatin, similarly to the distribution of HIF-2α (**Figure 2C**). Loss of HIF1β, which impedes HIF-2α binding to chromatin, resulted in a reduction of both HIF-2α and SET1B in the chromatin fraction, suggesting that SET1B binding to chromatin in ccRCC is partially dependent on HIF-2α (**Figure 2C**). However, SET1B deficiency did not prevent the chromatin localisation of HIF-2α, consistent with our prior observations with HIF-1α^24^.

To further determine if SET1B loss altered HIF-2α binding to HIF target genes, we performed ChIP-PCR for HIF-2α in wild-type, SET1B KO, or HIF-2α KO 786-0 cells, using primers targeting the promoters of *VEGFA* and *ANGPTL4*. Loss of SET1B did not affect HIF-2α binding to target gene loci, confirming that SET1B-dependent regulation of HIF-2α activity occurs at the chromatin level (**Figure 2D**). Both SET1B’s methyltransferase activity and binding to RNA Pol II appeared to be important for the regulation of HIF target genes, as loss of either SET1B or HIF-2α reduced H3K4me3 levels at the *VEGFA* promoter (**Figure S2E**) and resulted in reduced RNA Pol II abundance at exon 3 of *VEGFA* (**Figure 2E**). These results highlight the requirement for SET1B in driving transcription at HIF-2α target genes, likely dependent on both methyltransferase activity and RNA Pol II recruitment.

### SET1B facilitates HIF-2 driven ccRCC progression

To investigate the functional role of SET1B in ccRCC progression, we began by analysing the TCGA database https://www.cancer.gov/tcga. Specifically, we examined the mRNA expression levels of SET1B in the KIRC ccRCC dataset and assessed its correlation with patient survival (**Figure 3A**). Patients with the highest *SET1B* mRNA expression (top 25%) had a significantly lower survival rate than those with the lowest *SET1B* expression (**Figure 3A**). A positive correlation between *VEGFA* and *SET1B* mRNA expression was identified using cBioPortal (https://www.cbioportal.org/) (Figure **S3A**), and we also observed that SET1B loss led to reduced VEGF secretion in 786-0 cells (**Figure 3B**).

**Figure 3.**
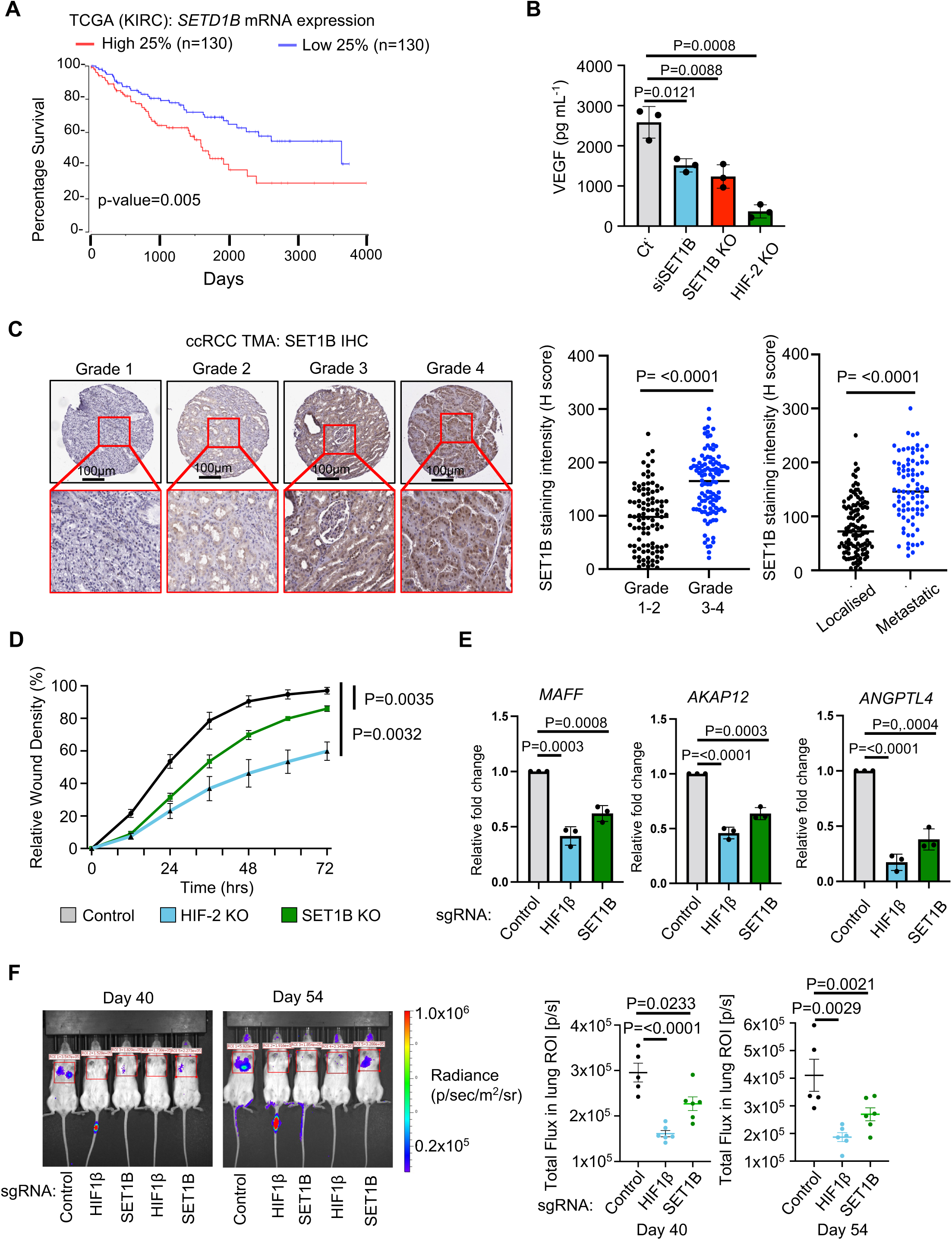
SET1B is required for sustained HIF-2-dependent activity in ccRCC. **(A)** High expression of (SET1B gene name) in kidney cancer is associated with poor outcomes. Kaplan-Meier survival analysis of TCGA data for ccRCC, comparing tumours in the highest and lowest quartiles of SETD1B mRNA expression. n=130 patients for each group. Log-rank test. **(B)** VEGFA ELISA in control, SET1B- depletion, SET1B KO and HIF-2α KO 786-0 cells. Cells were incubated for 48 hrs and before supernatants were collected. Graph is representative of three biological replicates. Two-way ANOVA. **(C)** Immunohistochemistry of SET1B was performed on a ccRCC tissue microarray. Samples were assessed and graded 1-4 with 1 being the least aggressive and grade 4 being the most aggressive. SET1B staining intensity across the tumour was calculated as an H score which accounts for the staining intensity and the % of positive cells detected. Samples were subdivided based on Grade and SET1B intensity was plotted. Significance was assessed using a two-way ANOVA. **(D)** Control, HIF-2α KO and SET1B KO 786-0 cells were embedded in Matrigel, and cellular invasion was measured over indicated time using the Incucyte. Graphs are representative for 3 biological replicates and depict the mean ± SD. Two-way ANOVA. **(E)** qPCR of HIF targets associated with metastasis (*MAFF*, *AKAP12* and *ANGPTL4*) in 786-0 cells depleted of HIF1β and SET1B using CRISPR (n = 3 biologically independent samples, mean ± SD). **(F)** ccRCC mouse xenograft model. Wildtype, HIF1β and SET1B depleted 786-0 cells expressing luciferase were injected into the tail vein of nude mice. Bioluminescence was measured and quantified from the lungs on Day 40 and Day 54 (Control =5, HIF1β KO = 6 and SET1B KO = 6). Mean ± SD. Two-way ANOVA.

To further investigate if SET1B is correlated with ccRCC progression, we examined SET1B levels using immunohistochemistry in a clinical cohort of ccRCC samples (n=317) (**Figure 3C**)^38^. SET1B levels were calculated using an H score (0-300), which encompasses the intensity of staining and the number of positive cells.

Increased SET1B levels within the tumour correlated with high tumour grade and in patients that had metastatic disease (**Figure 3C**).

To understand the functional requirement for SET1B in controlling HIF-2 activity in ccRCC, we focused on *in vitro* assays of metastatic potential and assessed the impact of SET1B loss on cell migration and invasion using a scratch and invasion assay. Depletion of either SET1B or treatment with the HIF-2 inhibitor (PT2385) in 786-0 cells had no effect on cell migration *in vitro* (**Figure S3B**). However, loss of HIF-2α reduced cellular invasion, confirming the role of HIF-2 in cellular invasion. Similar results were also observed for SET1B depletion, although not to the same extent as HIF-2α KO (**Figure 3D**). SET1B depletion also reduced the HIF-2 mediated transcription of several genes known to be determinants of metastases (*MAFF, AKAP12, ANGPTL4*) (**Figure 3E**)^39^, suggesting that SET1B coordinates cellular metastasis as well as angiogenesis in ccRCC.

Lastly, we explored the biological role of SET1B in ccRCC using an experimental *in vivo* metastasis model which measures the cell’s ability to extravasate and colonise the lung. SET1B and HIF1β were depleted in 786-0 luciferase-expressing cells using CRISPR, before the cells were injected into the tail vein of nude mice (**Figure S3C**). Cellular metastasis was determined by quantifying luciferase levels within the lung on days 40 and 54 (**Figure 3F**). Loss of HIF1β or SET1B resulted in very low, detection of lung luciferase signal (**Figure 3F**). Collectively, these findings indicate that SET1B helps drive angiogenesis and metastatic disease in a HIF-dependent manner.

### Depletion of SET1B potentiates HIF-2 inhibition in ccRCC

PT2385 (Belzutifan) is the first selective small-molecule inhibitor of HIF-2α activity in clinical use and inhibits the interaction between HIF-2α and HIF1β^40^ (**Figure S4A**). Given that PT2385 does not fully inhibit HIF-2 activity, we hypothesised that depleting SET1B could enhance PT2385-mediated suppression of HIF-2 signalling. Consistent with this notion, immunoprecipitation studies in 786-0 cells, treated and untreated with PT2385, revealed that while PT2385 disrupts the interaction between HIF1β and HIF-2α, it does not affect the interaction between HIF-2α and SET1B (**Figure 4A**), nor alter the protein levels of SET1B (**Figure S4B**). Moreover, cellular fractionation analyses revealed that PT2385 did not fully prevent HIF-2 or SET1B from binding chromatin (**Figure 4B**), consistent with some HIF-2 transcriptional activity remaining following PT2385 treatment. Therefore, to test if SET1B loss could enhance the activity of HIF-2 inhibition, we used siRNA-mediated SET1B depletion followed by treatment with different concentrations of PT2385. SET1B deficiency enhanced suppression of HIF-2-mediated transcription (**Figure 4C**). This combinatorial effect was also observed using SET1B shRNA (**Figure S4C**).

**Figure 4.**
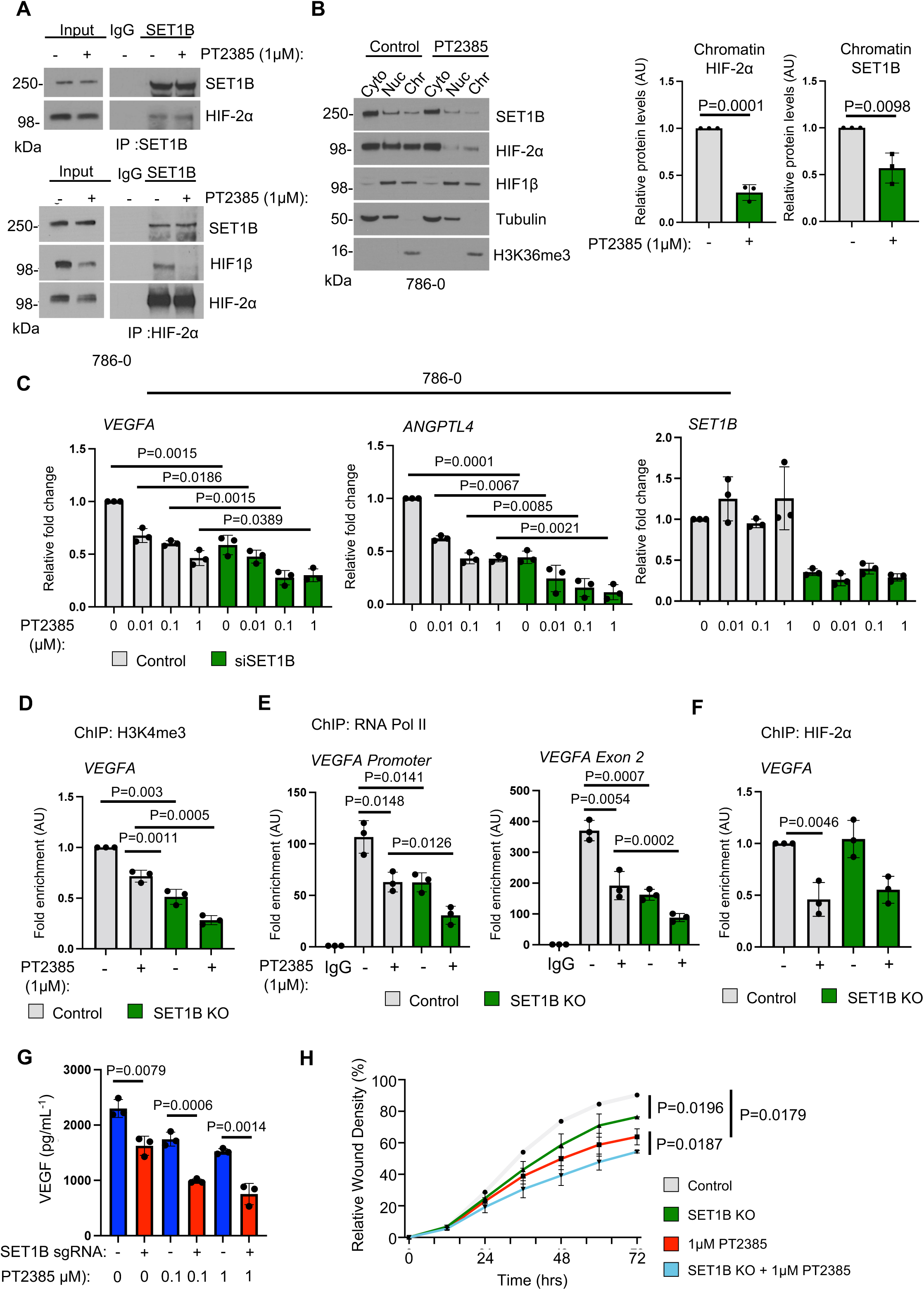
Depletion of SET1B potentiates HIF-2 inhibition in ccRCC. **(A)** Wildtype 786-0 cells were treated with and without 1 μM PT2385 for 24hrs. Endogenous SET1B and HIF-2α were immunoprecipitated and their interaction was evaluated by immunoblotting using the indicated antibodies. Immunoblots are representative of 3 biological replicates. **(B)** 786-0 cells were treated with and without 1 μM PT2385 were fractionated into cytoplasmic, nucleoplasm and chromatin fractions and SET1B and HIF-2α were monitored using immunoblotting. The ratio of SET1B and HIF-2α in the chromatin fraction was quantified using ImageJ (*n* = 3 biologically independent experiments, mean ± SD). **(C)** Wildtype 786-0 cells were transfected with a control and a SET1B siRNA for 48 hrs. 24 hrs prior to harvesting cells were treated with indicated concentrations of PT2385. HIF-2α activity was determined by performing qPCR of *VEGFA* and *ANGPTL4* (*n* = 3 biologically independent experiments, mean ± SD). SET1B mRNA levels was analysed to validate successful depletion. ChIP-PCR of H3K4me3 **(D)**, RNA Pol II **(E)** and HIF-2α **(F)** in wildtype and SET1B KO 786-0 cells treated in the presence and absence of 1 μM PT2385. ChIP-PCR was performed using primers surrounding the *VEGFA* promoter. (n= 3 biological replicates). Graphs show mean ± SD. Two-way ANOVA. **(G)** VEGFA ELISA in control and SET1B-depleted 786-0 cells. 24 hrs before collecting supernatants cells were treated with 1uM PT2385. Graph is representative of three biological replicates. Two-way ANOVA. **(H)** Control and SET1B KO 786-0 cells treated with and without 1 μM PT2385 were embedded in Matrigel, and cellular invasion was measured over indicated time using the Incucyte. Graphs are representative for 3 biological replicates and depict the mean ± SD. Two-way ANOVA.

Importantly, the combination of SET1B depletion with PT2385 enabled the use of lower PT2385 doses, leading to a more pronounced downregulation of HIF-2 activity (**Figure 4C**). A similar additive effect was observed in 769-P cells (**Figure S4D**), In contrast, RCC4 and RCC10 cells, which express both HIF-1α and HIF-2α, exhibited reduced HIF activity with SET1B depletion alone, without additional benefit from PT2385, suggesting potential compensatory mechanisms played by HIF-1α in these cells.

Mechanistically, concurrent SET1B loss and HIF-2α inhibition markedly reduced H3K4me3 levels at the promoter coupled with reduced binding of RNA Pol II levels at both the promoter and exon 2 of *VEGFA*, surpassing the effects observed with SET1B knockout or PT2385 treatment alone (**Figure 4D, E**). Notably, SET1B depletion did not alter HIF-2α binding in isolation or in combination with HIF-2 inhibition, consistent with SET1B modulating residual HIF-2α transcriptional activity on the chromatin (**Figure 4F**).

To further establish the functional significance of the synergy between SET1B depletion and HIF-2α inhibition, we evaluated VEGF secretion and cellular invasion in SET1B-depleted 786-0 cells following HIF-2 inhibition. PT2385 treatment significantly reduced VEGF secretion, with the combination of SET1B depletion and PT2385 resulting in a more substantial reduction compared to control cells (**Figure 4G**). This additive effect was similarly observed in cellular invasion assays, where SET1B loss alone decreased cellular invasion, but the combination with PT2385 had an enhanced inhibitory effect (**Figure 4H**).

### Targeting SET1B to circumvent HIF-2 inhibitor resistance

Recent clinical studies have identified that patients develop resistance to prolonged HIF-2α inhibition, which has been linked to increased p53 activity and the emergence of the G323E mutation within the HIF-2α inhibitor binding pocket^41^. The G323E mutation enables HIF-2α to re-associate with HIF1β, thereby promoting ccRCC progression. Targeting SET1B may therefore provide a strategy for combating HIF- 2α inhibitor resistance. To explore this, we reconstituted HIF-2α knockout 786-0 cells with either wild-type HIF-2α or the HIF-2α G323E mutant (**Figure 5A**) and examined the effects of SET1B depletion.

**Figure 5.**
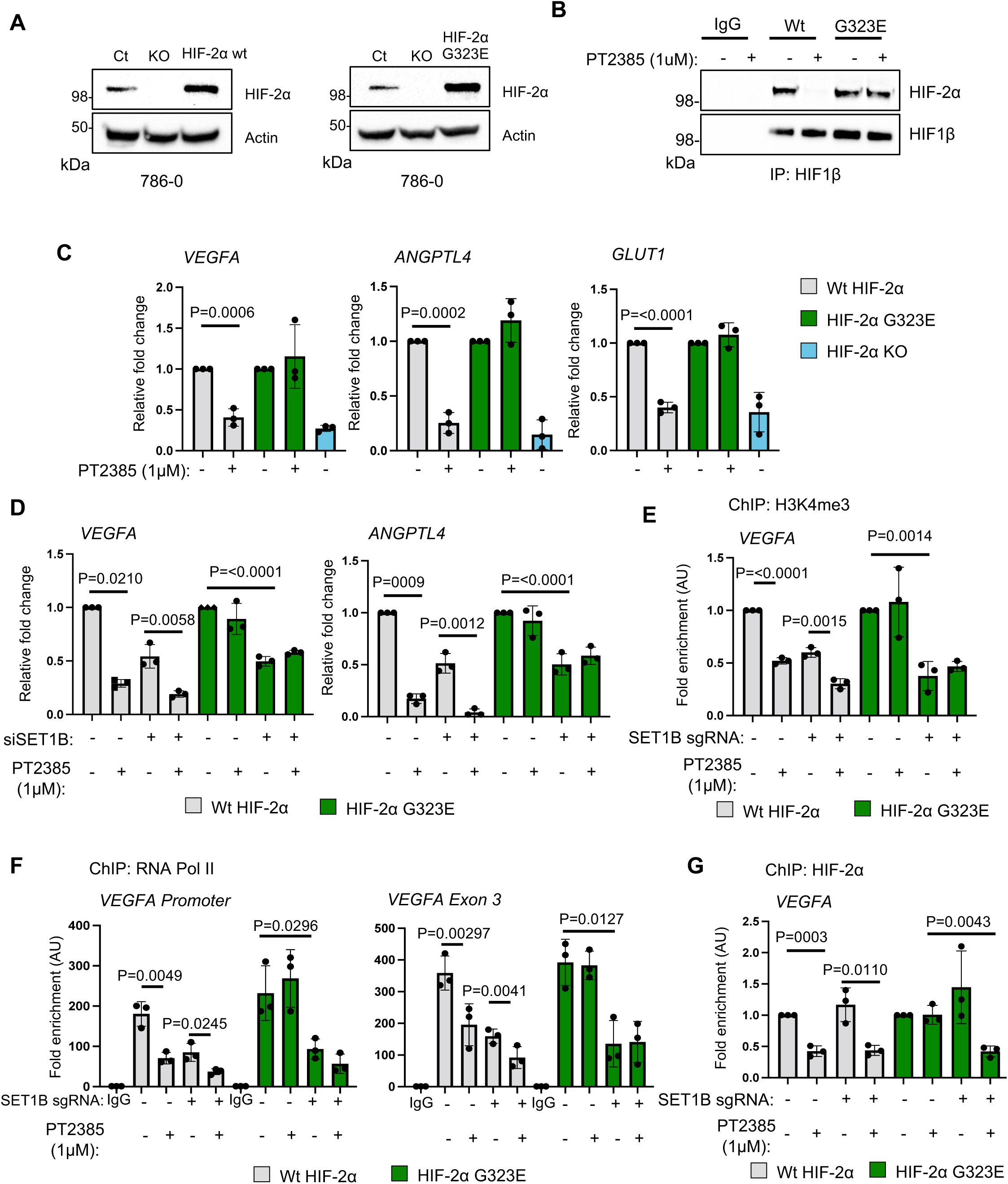
SET1B depletion overcomes resistance to HIF-2 inhibition in ccRCC. **(A)** Immunoblot of wildtype, HIF-2α KO and HIF-2α KO 786-0 cells reconstituted with wildtype HIF-2α or the HIF-2α G323E mutation. **(B)** HIF-2α KO cells reconstituted with wildtype and HIF-2 G323E mutation were treated with 1 μM PT2385 for 24 hrs. Endogenous HIF1β was immunoprecipitated and the interaction with HIF-2α was assessed by immunoblotting. Immunoblot is representative of 3 independent experiments. **(C)** HIF-2α KO cells reconstituted with wildtype and HIF-2 G323E mutation were treated with 1 μM PT2385 for 24 hrs. HIF-2α target genes and *SET1B* mRNA expression was assessed by qPCR (n= 3 biological replicates). Graphs show mean ± SD. Two-way ANOVA. **(D)** HIF-2α KO cells reconstituted with wildtype and HIF-2α G323E mutation were transfected with a SET1B siRNA for 48 hrs. 24 hrs for harvesting cells with treated with and without 1 μM PT2385. qPCR analysis was performed using primers targeting the *VEGFA* and *ANGPTL4* mRNA (n= 3 biological replicates). Graphs show mean ± SD. Two-way ANOVA. **(E)** ChIP of H3K4me3 and HIF-2α **(F)** in HIF-2α KO cells reconstituted with wildtype and HIF-2α G323E mutation following CRISPR-mediated SET1B depletion 24 hrs before harvesting cells were treated with 1 μM PT2385. ChIP-PCR was performed using primers targeting the *VEGFA* promoter (n= 3 biological replicates). Graphs show mean ± SD. Two-way ANOVA. **(G)** ChIP of RNA Pol II in HIF-2α KO cells reconstituted with wildtype and HIF-2α G323E mutation following CRISPR-mediated SET1B depletion 24hrs before harvesting cells were treated with 1 μM PT2385. ChIP-PCR was performed using primers targeting the *VEGFA* promoter and exon 3 (n= 3 biological replicates). Two-way ANOVA.

Reconstitution with the HIF-2α G323E mutant enabled HIF-2α to re-associate with HIF1β, even in the presence of PT2385 (**Figure 5B**), preventing the reduction in HIF- 2 target gene expression (**Figure 5C**). However, while 786-0 cells expressing the G323E mutant were resistant to PT2385, HIF-target gene transcription was still reduced following SET1B depletion (**Figure 5D**). SET1B depletion led to a decrease in H3K4me3 levels and RNA Pol II levels across the *VEGFA* gene in both wild-type and G323E mutant cells (**Figure 5E/5F),** without affecting HIF-2α binding (**Figure 5G**), indicating that SET1B loss reduced the transcriptional activity of the gene.

Taken together, these findings suggest that SET1B could serve as a potential therapeutic target to overcome HIF-2 inhibitor resistance and ccRCC progression, by blocking selective HIF-2 dependent transcription at the chromatin level (**Figure 6**).

**Figure 6.**
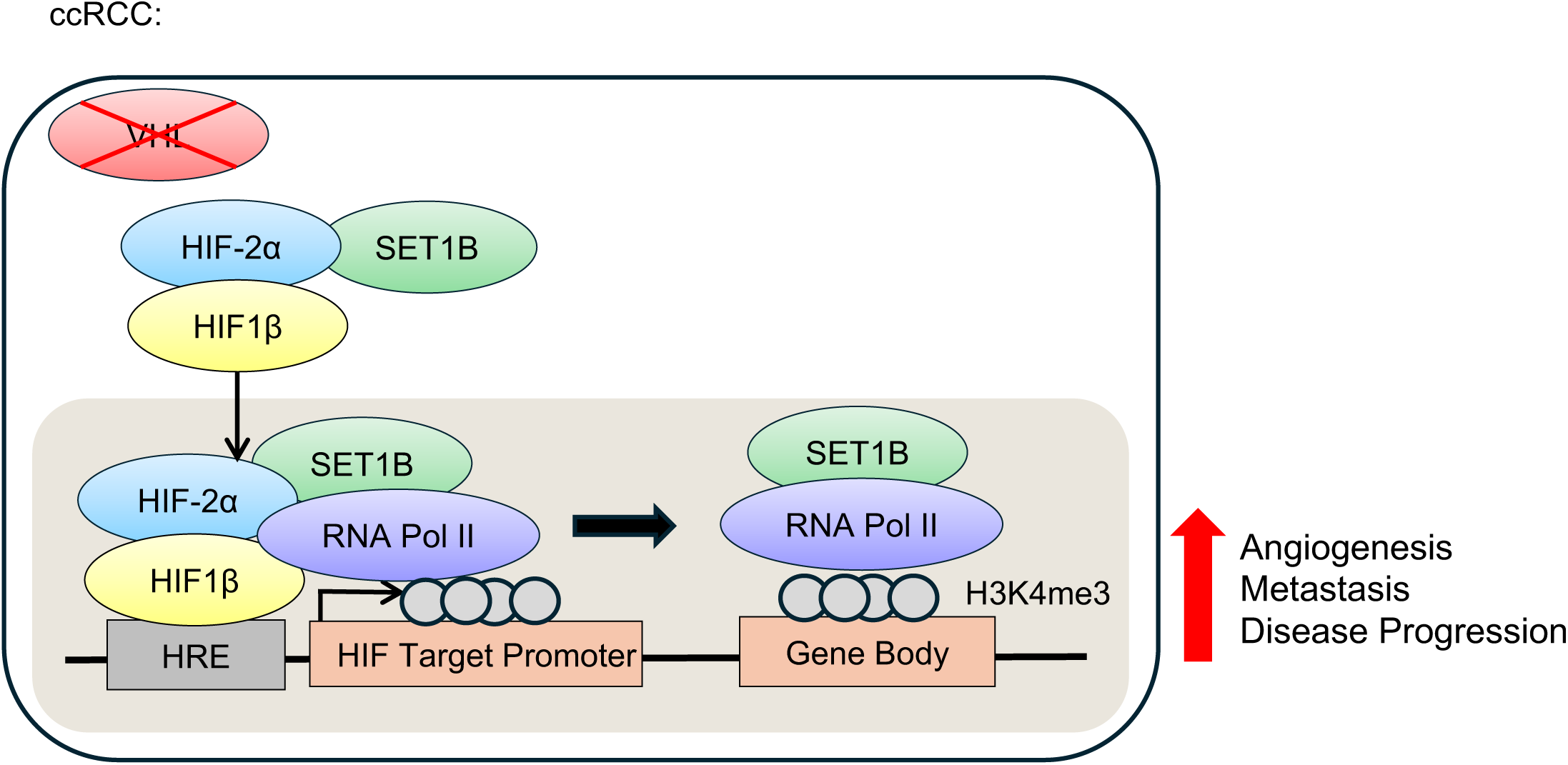
Model for the role of SET1B in ccRCC In ccRCC, Hypoxia-Inducible Factor 2 (HIF-2) is constitutively active due to the loss of von Hippel-Lindau (VHL) protein. SET1B is recruited to HIF target genes by the HIF complex, where it plays a critical role in initiating and sustaining HIF transcription. This is achieved through its histone H3 lysine 4 trimethylation (H3K4me3) activity and its interaction with the RNA Pol II complex. The enhanced HIF-2 transcription leads to increased angiogenesis, metastasis, and disease progression.

## Discussion

Although we initially hypothesised that SET1B primarily modulates H3K4me3 levels at HIF target loci, our findings reveal a more intricate mechanism of SET1B- dependent transcriptional regulation. Using a model of sustained HIF activation, we show that SET1B is not only essential for initiating HIF activity but also crucial for maintaining it over time. Notably, we present evidence that SET1B, in addition to its recruitment to gene promoters, binds across the gene bodies of HIF targets, with this binding closely associated with RNA Pol II occupancy.

Several studies have emphasised that H3K4me3 regulates RNA Pol II activity by facilitating the release of RNA Pol II from proximal-pause and promoting RNA Pol II elongation activity^25,32^. However, as SET1B appears to travel with the RNA Pol II complex along the DNA, it is possible that SET1B may have additional roles in transcriptional regulation. This notion would be consistent with other studies focusing on the regulation of HIF target genes, where transcriptional complexes such as DNAPK, FACT, TIP60, and ZMYND8 enhance HIF-dependent transcription by modulating RNA Pol II activity^15,17,18,42^. It will be helpful in future work to determine whether the spreading of H3K4me3 into gene bodies contributes to the regulation of RNA Pol II activity at HIF target genes, as observed for the establishment of cellular identity in embryonic stem cells^43^.

Our studies on SET1 functional domains show that both the DPR motif, as well as the RMM and SET domains are required for modulating HIF transcriptional activity. The involvement of the DPR motif of SET1B is consistent with prior studies, and the DPR motif has been shown to modulate transcription through binding to WDR82 in mouse embryonic stem cells^32^. While RRMs are known to bind RNA, both SET1A and SET1B each contain a single RRM, in contrast to other RNA-interacting proteins that generally require two RRMs for RNA binding^44^. As loss of the RRM reduces the ability of SET1B to interact with RNA Pol II, this raises the possibility that SET1B may recognise specific RNA species or alternatively, RNA binding may modulate the methylase activity of SET1B which has been observed for other enzymes^45^. It will be of interest to address RNA binding by SET1B in future studies, but it is noteworthy that the yeast SET1 complex binds nascent RNA and this binding is required for the deposition of H3K4me3 ^46^.

The requirement of the SET domain for SET1B-mediated regulation of HIF target gene was evident in our studies, but the role of H3K4me3 in transcription remains debated. Recent studies have highlighted the importance of H3K4me3 in RNA Pol II pausing release at the promoter^25^, and our data suggests that SET1B methylase activity may contribute to transcriptional changes even with modest changes in H3K4me3 levels. However, it is also possible that SET1B may methylate other proteins aside from histones, just as SET1A monomethylates YAP, promoting YAP activity and preventing its nuclear export^47^. Similarly, studies have emphasised the importance of RNA Pol II lysine methylation in transcription regulation^48,49^.

In ccRCC, loss of VHL leads to constitutive HIF activation, with several studies highlighting the importance of HIF-2α in promoting angiogenesis and metastasis^50–52^. Using a panel of ccRCC-derived cell lines, we found that SET1B helps to sustain HIF-dependent activity at the chromatin level. Additionally, SET1B expression correlates with disease grade and metastasis in patient samples, with particularly high levels observed in metastatic ccRCC. Furthermore, depletion of SET1B impaired the ability of ccRCC cells to survive *in vivo* in an experimental metastasis model. These findings underscore the role of SET1B in sustaining HIF-2α activity and driving the expression of genes involved in angiogenesis and metastatic spread.

The development of HIF-2α-specific inhibitors are a promising therapeutic approach for ccRCC, now in routine clinical practice in some nations, but HIF-2α inhibition can have off-target effects due to its roles in other physiological processes, including immune cell function and carotid body activity^53–55^. Pursuing selective modulation of SET1B activity may be a potential strategy for target ccRCC while reducing the impact on other essential HIF-2-dependent functions. Finally, resistance to HIF-2α inhibitors has been observed in some patients after prolonged treatment with Belzutifan. Several mechanisms have been proposed, including the upregulation of p53 activity or the G323E mutation within the inhibitor binding pocket^41^. Our findings now indicate that targeting SET1B in patients with the G323E mutation could be a viable strategy to reduce HIF-2-driven pathways involved in angiogenesis and metastasis. However, whether therapeutic targeting of SET1B is possible remains to be seen. Inhibitors for the SET family of methyltransferases are being actively pursued^56–58^, but our studies also indicate that inhibition of SET activity may not be sufficient, and that other domains, including those involved in the recruitment of Pol II and RNA binding need to be considered.

## Methods

### Cell lines and reagents

786-0 wildtype and 769-P cells were maintained in RPMI-1640 (Sigma) supplemented with 10% foetal calf serum (FCS). HeLa, A549, HEK293T, RCC4 and RCC10 cells were maintained in DMEM (Sigma) supplemented with 10% FCS. Cells were maintained in a 5% CO2 incubator at 37 °C. PT2385 was acquired from Biovision (B1920) and was added to cell cultures at indicated concentrations.

### Plasmids

Plasmids used: pKLV-U6gRNA-EF(BbsI)-PGKpuro2ABFP (Addgene plasmid, catalog no. 62348), LentiCRISPRv2 (sgRNA/Cas9, F. Zhang Addgene catalog no. 52961), pHRSIN-pSFFV-HA-pPGK-Puro, pHRSIN-pSFFV-pPGK-Puro, pMD.G (Lentiviral VSVG), pMD.GagPol (Lentiviral Gag/Pol). HIF-2 truncation mutant constructs were generated from pCDNA3 HIF-2α and cloned into the pHRSIN-pSFFV backbone with puromycin resistance using NEBuilder HiFi (NEB). pCDNA3 HIF-2 was cloned into pHRSIN-pSFFV backbone and Gibson cloning was used to generate the G323E mutant. Primers for cloning the G323E and truncation mutants are shown in Supplementary Table 1. The SET1B construct was a gift from D. Skalnik’s laboratory. SET1B was cloned into pHRSIN-pSFFV backbone with puromycin resistance using NEBuilder HiFi (NEB) as described previously ^24^.

Primers for HIF-2a cloning are shown in Supplementary Table 1.

### Lentiviral production and transduction

Lentivirus was produced by transfection of HEK293T cells using Fugene (Promega) at 70–80% confluency in six-well plates, with the appropriate pHRSIN vector and the packaging vectors pCMVR8.91 (gag/pol) and pMD.G (VSVG). Viral supernatant was harvested at 48 h, filtered (0.45-μm filter), and stored at −80 °C. For transduction, cells were seeded on 24-well plates in 500 µl medium, 500 µl viral supernatant added and plates centrifuged at 1,800 r.p.m., 37 °C for 1 h. Antibiotic selection was applied from 48 h.

### CRISPR–Cas9 targeted deletions

Gene-specific CRISPR sgRNA sequences were taken from the TKO library or designed using E-CRISP (http://www.e-crisp.org/E-CRISP/), with 5′CACC and

3′CAAA overhangs, respectively. SgRNAs were ligated into the LentiCRISPRv2 or pKLV-U6gRNA(BbsI)-PGKpuro2ABFP vector and lentivirus produced as described. Transduced cells were selected with puromycin and were generally cultured for 9–10 days before subsequent experiments to allow sufficient times for depletion of the target protein. KO clones were isolated from the sgRNA-targeted populations by serial dilution or FACS. SgRNAs used are shown in Supplementary Table 1.

### Immunoblotting

Cells were lysed in an SDS lysis buffer (1% SDS, 50 mM Tris (pH 7.4), 150 mM NaCl, 10% glycerol and 5 µl ml^−1^ Benzonase (Sigma)) for 10 min before heating at 90 °C for 5 min. Proteins were separated by SDS–PAGE, transferred to polyvinylidenedifluoride (PVDF) membranes, probed with appropriate primary and secondary antibodies and developed using enhanced chemiluminescent (ECL) or Super signal West Pico Plus Chemiluminescent substrate (Thermo Scientific).

### Quantitative PCR

Total RNA was extracted using the RNeasy Plus minikit (Qiagen) following the manufacturer’s instructions and then reversed transcribed using Protoscript II Reverse Transcriptase (NEB). Template cDNA (20 ng) was amplified using the ABI 7900HT Real-Time PCR system (Applied Biotechnology or Quantstudio 7, Thermo Scientific) reactions Transcript levels of genes were normalized to a reference index of a housekeeping gene (β-actin). Primers sequences are shown in Supplementary Table 1.

### Immunoprecipitation

786-0 cells were lysed in 1% Triton TBS, with 1x Roche complete EDTA-free protease inhibitor cocktail for 30 min at 4 °C. Lysates were centrifuged at 14,000 r.p.m. for 10 min, supernatants collected and then diluted to 0.1% detergent for preclearing with Protein G magnetic beads (Thermo Scientific) for 2 h at 4 °C. Supernatants were then incubated with primary antibody overnight (rotation at 4 °C). Protein G magnetic beads were then added for 2 h, and samples were then washed three times. Bound proteins were eluted in 2× SDS loading buffer, separated by SDS–PAGE and immunoblotted.

### Subcellular fractionation

A total of 10 × 10^6^ 786-0 cells were washed in PBS, lysed in Buffer A (10 mM HEPES, 1.5 mM MgCl_2_, 10 mM KCl, 0.5 mM DTT and EDTA-free protease inhibitor cocktail tablet, Roche) and incubated with rotation at 4 °C for 10 min. Supernatant containing cytosolic fractions were collected by centrifugation (1,400g for 4 min at 4 °C). The nuclear pellet was resuspended in Buffer B (20 mM HEPES, 1.5 mM MgCl_2_, 300 mM NaCl, 0.5 mM DTT, 25% glycerol, 0.2 mM EDTA and EDTA-free protease inhibitor cocktail tablet) for 10 min on ice to separate nucleoplasmic and chromatin fractions. Samples were centrifuged at 1,700g for 4 min at 4 °C, separating the soluble nucleoplasm from the insoluble chromatin fraction. The chromatin fraction was solubilized in 2× SDS loading buffer containing 1:500 Benzonase (Sigma).

### VEGF ELISA

Cells were plated in six-well plates and treated with 1% or 21% O_2_ for 24 h. Culture supernatants were collected, centrifuged at 1,500 r.p.m. for 10 min at 4 °C and analyzed using the Human VEGF Quantikine ELISA kit (R&D Systems) according to the manufacturer’s instructions.

### ChIP–qPCR

786-0 and HeLa cells grown on 15-cm dishes up a 2 × 10^6^ density and treated with 1% formaldehyde for 10 min to crosslink proteins to chromatin. The reaction was quenched with glycine (0.125 M for 10 min at room temperature). Cells were then washed in ice-cold PBS twice, scraped in tubes, and centrifuged at 800 r.p.m. for 10 min before lysis in 500 µl of ChIP lysis buffer (50 mM Tris-HCl (pH 8.1), 1% SDS, 10 mM EDTA, Complete Mini EDTA-free protease inhibitor). Samples were incubated on ice for 10 min and diluted 1:1 with ChIP dilution buffer (20 mM Tris-HCl (pH 8.1), 1% (v/v) Triton X-100, 2 mM EDTA and 150 mM NaCl). Samples were then sonicated in tubes and beads for 20 cycles of 30 s on and 30 s off in a Biorupter (Diagenode), followed by centrifugation for 10 min at 13,000 r.p.m. at 4 °C. Supernatants were collected and 20 µl stored at −20 °C as the input sample. A 200 µl aliquot of the remaining sample was diluted with ChIP dilution buffer to 1 ml and precleared using 20 µl Protein G magnetic beads (4 °C, 2 h, rotating); 1 ml of sample was immunoprecipitated with the appropriate primary antibody (4 °C, overnight, rotating). Protein G magnetic beads (25 μl) were added and incubated for a further 2 h at 4 °C. The beads were washed sequentially for 5 min each with wash buffer 1 (20 mM Tris-HCl (pH 8.1), 0.1% (w/v) SDS, 1% (v/v) Triton X-100, 2 mM EDTA, 150 mM NaCl), wash buffer 2 (Wash Buffer 1 with 500 mM NaCl), wash buffer 3 (10 mM Tris-HCl (pH 8.1), 0.25 M LiCl, 7 1% (v/v) NP-40, 1% (w/v) Na-deoxycholate and 1 mM EDTA), and twice with TE buffer (10 mM Tris-HCl (pH 8.0) and 1 mM EDTA). Bound complexes were eluted with 120 μl elution buffer (1% (w/v) SDS and 0.1 M Na-bicarbonate), and crosslinking reversed by addition of 0.2 M NaCl, and incubation at 65 °C overnight with agitation (300 r.p.m.). Protein was digested with 20 μg proteinase K (Thermo Scientific) for 4 h at 45 °C. RNase H (Thermo Scientific) was added for 30 min at 37 °C and DNA purified using the DNA minielute kit (Qiagen). DNA underwent qPCR analysis, and results expressed relative to input material. Primers sequences are shown in Supplementary Table 1.

### shRNA-mediated RNA depletion

Oligonucleotides were designed with a TTCAAGAGA hairpin, using shRNA sequences from the Broad Institute RNAi Consortium shRNA Library (Table S5). Sequences were cloned into pC.SIREN.Puro by digestion and ligation. Cells were transduced with the indicated vector or a scrambled shRNA control, and assays performed after at least seven days, following Puromycin selection. The shRNA constructs were purchased from Sigma and cloned into the pSIREN vector. The clone IDs for SET1B are TRCN0000237963 (sh*SET1B*-#1)

### siRNA-mediated depletion

Cells were transfected with siRNA SMARTpools for *SETD1B* (Dharmacon), SETD1B UTR siRNA (Dharmacon), Universal Negative Control using Lipofectamine RNAi MAX (Thermo Fisher). Cells were harvested after 48 h for further analysis by flow cytometry, qPCR or immunoblot.

### Analysis of TCGA expression and survival data

SETD1B mRNA expression and survival data for ccRCC tumours was obtained from The Cancer Genome Atlas (TCGA). The R package survminer was used to perform a log-rank test and plot a Kaplan-Meier curve.

### Migration assay

1.2x10^4^ 786-0 cells were seeded per well in 96-well ImageLockTM plates (4379, Essen BioScience) and incubated for 24 h. A minimum of 18 wells per condition were seeded in each biological replicate. If the effect of PT2385 was tested, the compound was added to the wells once the cells were attached. The WoundMakerTM (4563, Essen BioScience) is a 96-spin mechanical device designed to create 700-800 μm wide homogeneous wounds in cell monolayers on 96-well ImageLockTM plates. Prior to use after storage, the WoundMakerTM lid was washed with sterile distilled water for 5 min and with 70% ethanol for another 5 min. After washing, the scratch was performed to create precise wounds in all wells of the ImageLockTM plate. After wounding, cells were washed with fresh media to prevent dislodged cells from settling and reattaching. When PT2385 was tested, the new media contained 1 μM PT2385. Once the wound was made, plates were placed into the IncuCyte ZOOM® for 24 h in the case of 786-0 cells or 72 h in the case of RCC4 cells. Scanning was performed using a 10x objective and scheduled every 2 hours. Migration ability of different cells and conditions was analysed through two integrated metrics that the IncuCyteTM Software calculates based on the processed images: wound width and wound confluence. Wound width represents the average distance (μm) between the leading edge of the population of migrating cells (scratch wound mask) within an image. Wound confluence determines the percentage of wound area that is occupied by cells, and it relies on the initial scratch wound mask to differentiate the wounded from the non-wounded region. Debris or other matter within the wound is quantified as confluence and is not background subtracted.

Therefore, the wound confluence values at the initial time point may not be 0%.

### Clinical Samples

All patient tissue samples were used in accordance with approval granted by the Northumberland, Tyne and Wear NHS Strategic Health Authority Research Ethics Committee (reference 2003/11; The Freeman Hospital) and informed consent from all patients.

### Immunohistochemistry analysis

Immunohistochemistry was performed using tissue microarrays containing 0.6-mm cores of renal cancer and control kidney tissue as described^38^. Sections were immunostained with anti-SET1B 1:100 (Atlas Antibodies HPA021667) and viewed using Aperio CS2 (Leica Biosystems). SET1B intensity was compared across normal vs tumour tissue and across the different clinical stages of ccRCC which were graded by two independent clinicians who were blinded, and the H score was calculated.

### Cell invasion assay

96-well ImageLockTM plate wells were coated with a thin layer of Matrigel® Growth Factor Reduced Basement Membrane Matrix (354230, Corning®). To this end, Matrigel® stock solution was dissolved in cold cell culture media to a final concentration of 100 μg/mL. Wells were coated with 50μL of the solution and the plate was placed in a 37C incubator, 5% CO2 overnight. Matrigel® was removed and, either 1.2x10^4^ cells in the case of the 786-0 cell line or 2x10^4^ in the case of the RCC4 cell line were seeded per well and incubated for 24 h. In this case, a minimum of 10 wells per condition were used per biological replicate. When necessary, PT2385 was added to the wells once the cells were attached. After this time, the scratch was performed to create precise wounds in all wells of the ImageLockTM plate. Wells were washed once with cell culture media and 50 μL Matrigel® (8mg/mL) was added to each well carefully, avoiding the formation of bubbles. The plate was placed in the incubator for 30 min prior to the addition of 100 μL cell culture media containing PT2385 or not. The plate was then placed into the IncuCyte ZOOM® for 5 days. Scanning was performed using a 10x objective and scheduled every 4 h. The WoundMakerTM was washed before and after using as described in ‘Migration assay’. The invasion ability was analysed using the relative wound density (RWD). Like wound confluence, RWD also relies on the initial scratch wound mask to differentiate between cell-occupied and cell-free regions of the image, because it calculates the density of the wound region relative to the density of the cell region.

This parameter is the only recommended metric for cell invasion because 1) the mild texture of the ECM can hinder the ability of the analysis algorithm to apply an appropriate initial scratch wound mask and confluence mask and consequently result in a misleading wound confluence metric, and 2) cells invading through a 3D matrix usually exhibit an elongated phenotype with filopodia that extend into the ECM. Furthermore, one leader cell is generally followed by numerous other cells, forming tracts, as opposed to a leading edge of cells seen in a migration assay. For this reason, the scratch wound masks may not best represent the invading population and wound width measure can thus be inaccurate and misleading.

### Luciferase mouse metastasis and imaging

Cells were harvested and washed in Dulbecco’s phosphate buffer saline (D-PBS, without calcium and magnesium). Immunodeficient mice (NOD.Cg-*Prkdc^scid^ Il2rg^tm1Wjl^/SzJ*, RRID:IMSR_JAX:005557, purchased from Charles River UK) were housed at a density of 5 animals per individually ventilated cage with ad libitum access to food and water in a specific pathogen free unit. Mice were administered 5x10^5^ cells in D-PBS via intravenous injection to the lateral tail vein. Cells were administered in a blinded and randomised manner between cages. After 12 days in vivo bioluminescence imaging was performed with an IVIS imager (Perkin Elmer). Mice were administered 120 mg/kg D-Luciferin (Source Bioscience, prepared in D-PBS) via intraperitoneal injection and after five minutes anesthetised with isoflurane then placed in the IVIS for image acquisition (120 s image capture). After all images were collected regions of interest were drawn over the lungs and background subtracted radiance was determined. Images were captured every seven days until the experiment was terminated 75 days after injection. All experiments were performed according to protocols approved by the UK Home Office regulations, UK Animals (Scientific Procedures) Act of 1986 and were approved by the University of Cambridge Animal Welfare and Ethical Review Board.

### Statistical analyses

Quantification and data analysis of experiments are expressed as mean ± s.d. and *P* values were calculated using analysis of variance (ANOVA) or two-tailed Student’s *t*-test for pairwise comparisons and were calculated using Graphpad Prism v.8. Qualitative experiments were repeated independently to confirm accuracy.

## Acknowledgments

We thank all members of the Nathan and Ortmann groups for their helpful comments on the work and manuscript. The authors gratefully acknowledge the NIHR BRC flow cytometry facility, and the support of the Cambridge Institute of Medical Research Mass Spectrometry Facility. This work was funded by a Wellcome Senior Clinical Research Fellowship to JAN (215477/Z/19/Z), a Lister Institute Research Fellowship to JAN. BMO supported by Newcastle Academic Tract (NUAcT) Fellowship STR/0260/NACT/BO 02. AOS was supported by a package from the Wellcome Sanger Institute and Wellcome Trust. This work was also supported by a CRUK Cambridge Centre Urological Malignancies Programme Pump Priming award (Cancer Research UK Cambridge Centre [C9685/A25177 and CTRQQR- 2021\100012]). GDS is supported by The Mark Foundation for Cancer Research [RG95043], the Cancer Research UK Cambridge Centre [C9685/A25177 and CTRQQR-2021\100012] and NIHR Cambridge Biomedical Research Centre (NIHR203312). The views expressed are those of the author and not necessarily those of the NIHR or the Department of Health and Social Care.

## Author contributions

Conceptualization: BMO, JAN, Methodology: BMO, JAN, EA, LW, AH, SC, ALH, Investigation: BMO, JB, RS, EA, LW, AOS. Visualization: BMO, JB, EA, LW Supervision: BMO, JAN, CNR. Writing—original draft: BMO, JAN, Writing—review & editing: all authors

## Competing interests

GDS has received educational grants from Pfizer and AstraZeneca; consultancy fees from Evinova; travel expenses from MSD; he is Clinical lead (urology) National Kidney Cancer Audit and Topic Advisor for the NICE kidney cancer guideline. All other authors declare that they have no competing interests

---

**Figure S1.**
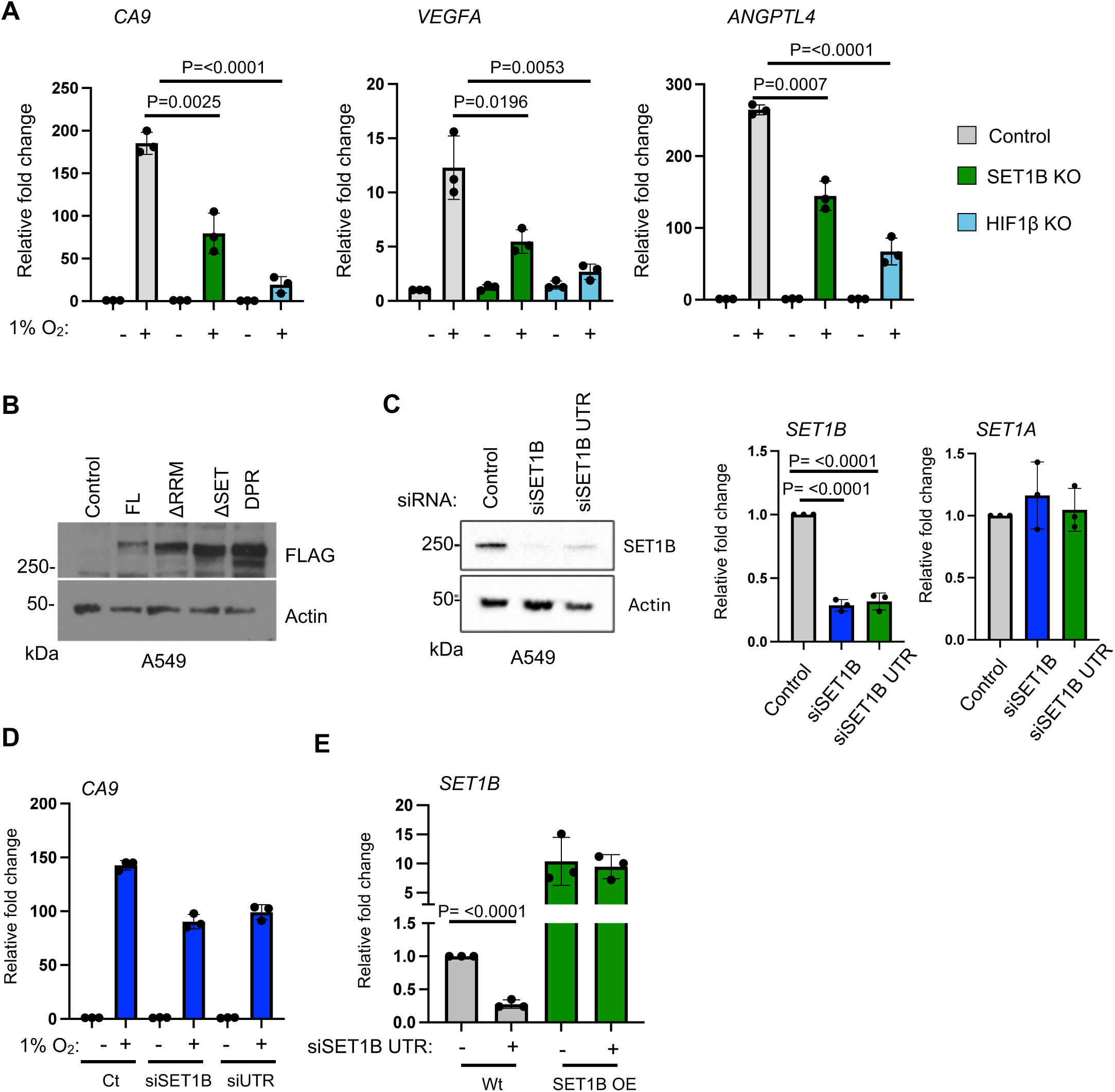
SET1B interacts with the RNA Pol II complex and requires multiple functional domains to coordinate HIF activity. **(A)** qPCR analysis of HIF target genes. A549 cells were transduced with sgRNAs targeting HIF1B and SET1B. Cells were incubated at 21% or 1% oxygen for 24 hours before qPCR was performed to analyse the mRNA expression levels of HIF target genes CA9, VEGFA and ANGPTL4 (n= 3 biological replicates). Graphs show mean ± SD. Two-way ANOVA. **(B)** Immunoblot analysis of A549 cells stably expressing full-length SET1B and various SET1B truncation mutants**. (C)** A549 cells were transfected with a pool of SET1B siRNAs and an siRNA targeting the 3’UTR of SET1B for 48 hrs. Samples were assessed using immunoblotting and qPCR using primers targeting *SET1B* and *SET1A*. Graphs are representative for 3 biological replicates and depict the mean ± SD. Two-way ANOVA. **(D)** A549 cells were transfected with a pool of SET1B siRNAs and an siRNA targeting the 3’UTR of SET1B for 48 hrs. 24 hrs before sample collection cells were incubated at 21% and 1% oxygen for 24 hrs. qPCR was performed using primers targeting the CA9 and SET1B. Graphs are representative for 3 biological replicates and depict the mean ± SD. Two-way ANOVA. **(E)** Wildtype and A549 cells stably with stable overexpression of FL SET1B were transfected with an siRNA targeting 3’UTR of SET1B for 48 hrs. qPCR was performed using primers targeting SET1B Graphs are representative for 3 biological replicates and depict the mean ± SD. Two-way ANOVA.

**Figure S2.**
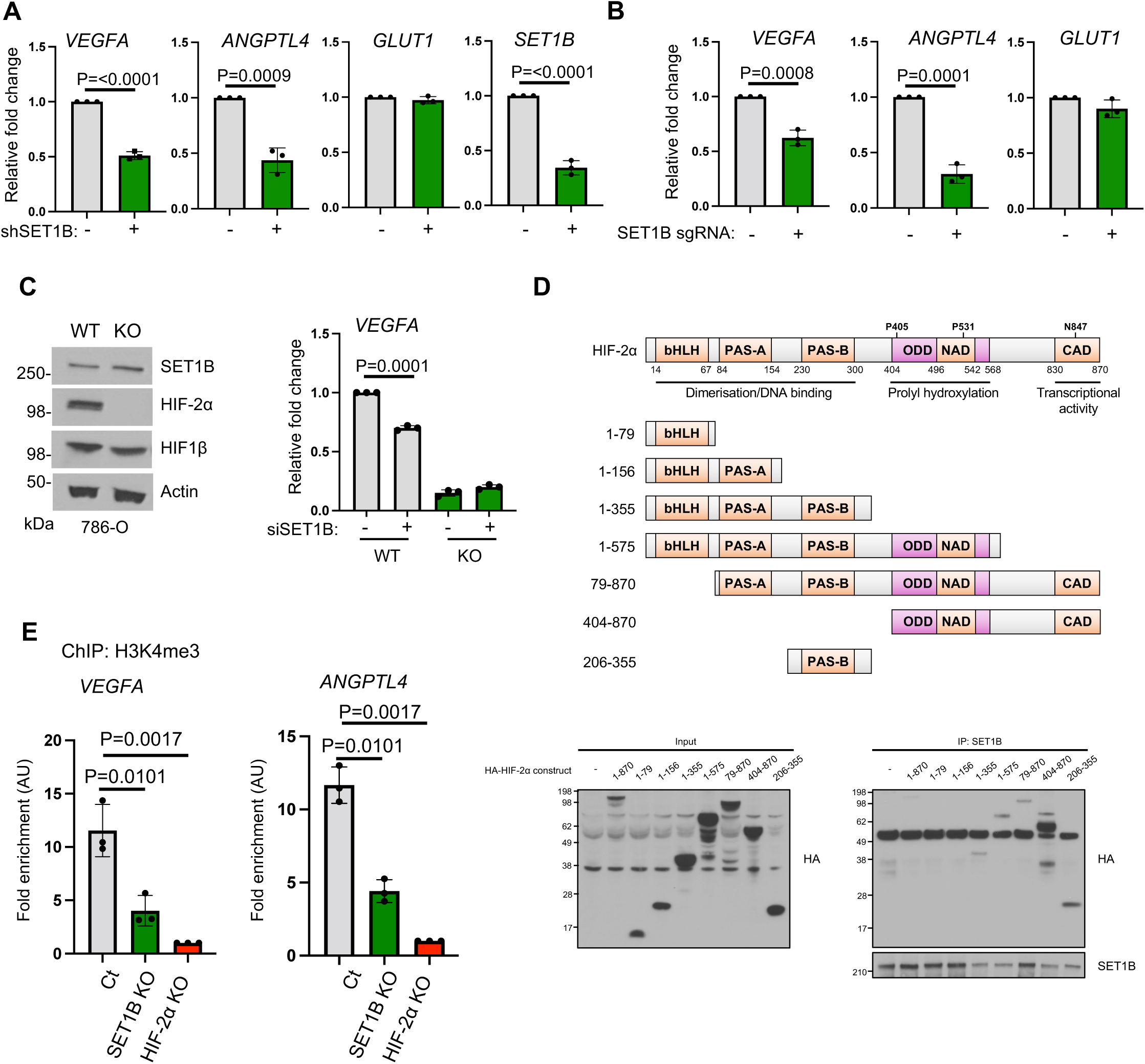
SET1B is required for sustained HIF-2 activity. 786-0 cells were transduced with an shRNA **(A)** or an sgRNA targeting SET1B **(B)** and the expression of HIF-2α target genes (*VEGFA, ANGPTL4, GLUT1*) and *SET1B* were assessed using qPCR. Graphs are representative for 3 biological replicates and depict the mean ± SD. Two-way ANOVA. **(C)** Immunoblot and qPCR of wildtype and HIF-2α KO 786-0 cells with indicated antibodies and qPCR was performed on *VEGFA*. Graphs are representative for 3 biological replicates and depict the mean ± SD. Two-way ANOVA. **(D)** Schematic diagram of full-length HIF-2α and truncation mutants. HEK293T cells were transfected with full length HIF-2α and the various truncation mutants using calcium phosphate for 48hrs. Cells were incubated for 6 hrs at 1% oxygen and endogenous SET1B was immunoprecipitated to assess their interaction with SET1B using immunoblotting. Immunoblot is representative of three biological replicates. **(E)** ChIP-PCR of H3K4me3 in wildtype, SET1B KO and HIF-2α KO 786-0 cells. ChIP-PCR was performed using primers targeting the promoters of *VEGFA* and *ANGPTL4.* Graphs are representative for 3 biological replicates and depict the mean ± SD. Two-way ANOVA.

**Figure S3.**
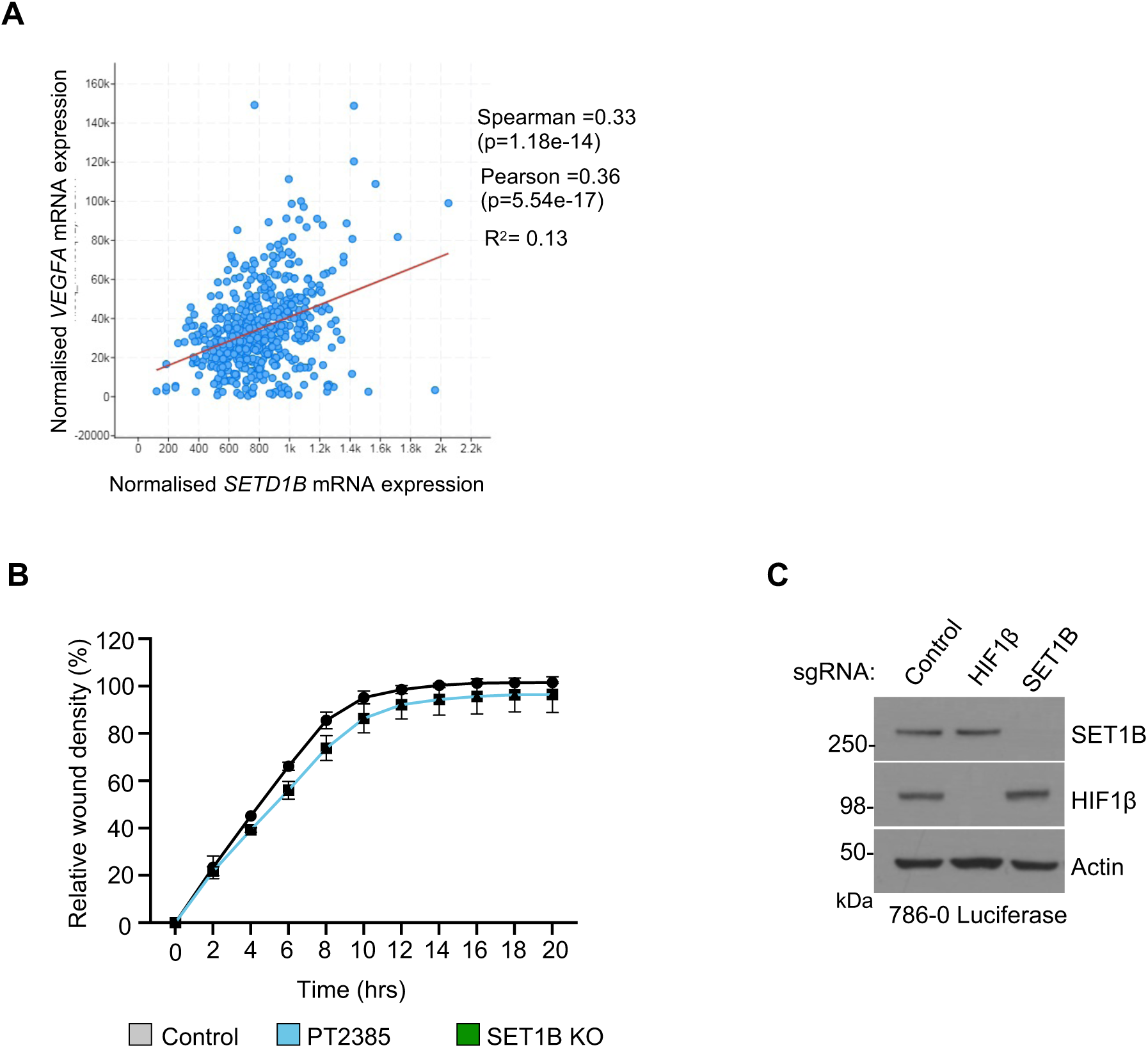
The effect of SET1B depletion on *VEGF* expression and on cell migration in ccRCC. **(A)** Comparison of the expression of *VEGFA* mRNA vs SETD1B in ccRCC using cBioportal. **(B)** Cell migration assay. Control SET1B KO and PT2385 treated cells were plated at high density before a scratch was made in the well. Cell migration was measured using the Incucyte at indicated time point. Graph is representative of 3 biological replicates. **(C)** Immunoblot of control sgRNA depleted HIF1β and SET1B 786-0 luciferase cells. Immunoblot was conducted before implantation into mice.

**Figure S4.**
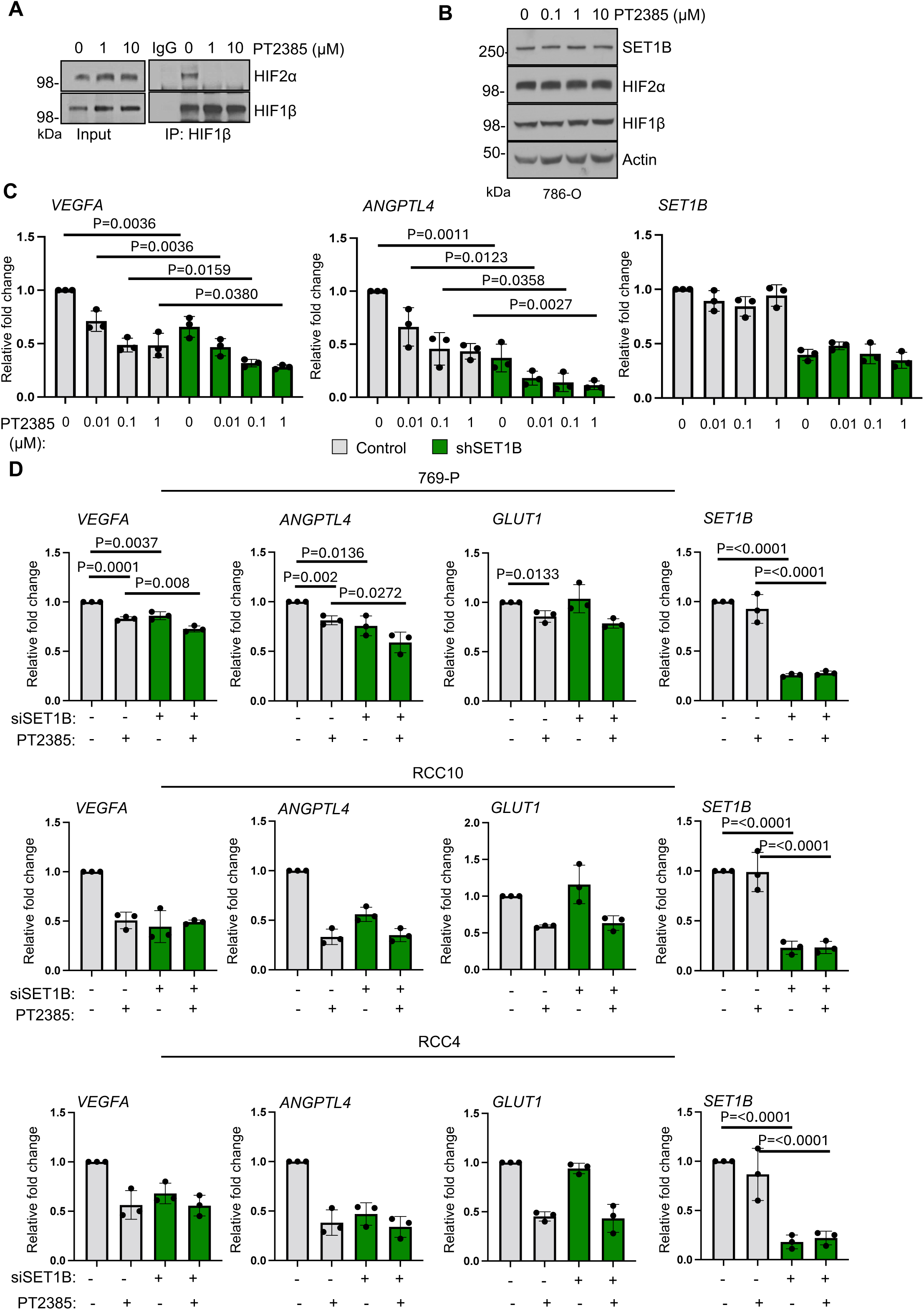
Loss of SET1B enhances the efficacy of HIF-2 Inhibition. **(A)** HIF1β was immunoprecipitated in 786-0 cells and its interaction with HIF-2α was assessed using immunoblotting following treatment indicated concentration of PT2385 for 24 hours. Immunoblot representative of three biological replicates. **(B)** Effect of PT2385 on expression of SET1B and HIF-2α, 786-0 cells were treated with indicated concentrations of PT2385 for 24 hrs. Immunoblot is representative of three biological replicates. **(C)** Wildtype 786-0 and 78-0 cells expressing an shRNA targeting SET1B were treated with indicated concentrations of PT2385 prior to harvesting. HIF-2α activity was determined by performing qPCR of *VEGFA* and *ANGPTL4* (*n* = 3 biologically independent experiments, mean ± SD). SET1B mRNA levels was analysed to validate successful depletion. **(D)** Wildtype 769-P, RCC4 and RCC10 cells were transfected with a control or a SET1B siRNA for 48 hrs. 24 hrs prior to harvesting cells were treated with indicated concentrations of PT2385. HIF- 2α activity was determined by performing qPCR of VEGFA and ANGPTL4 (*n* = 3 biologically independent experiments, mean ± SD).

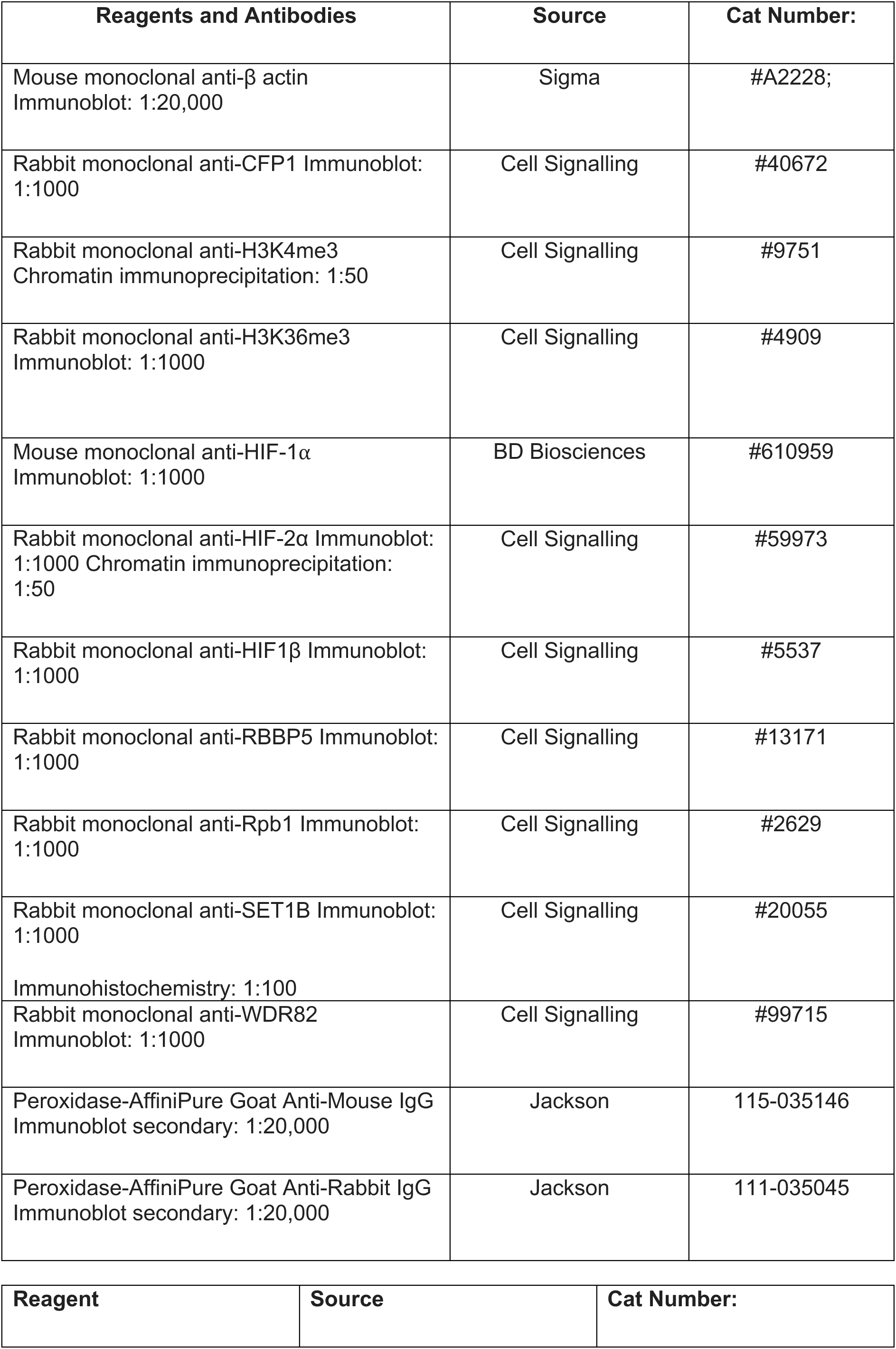
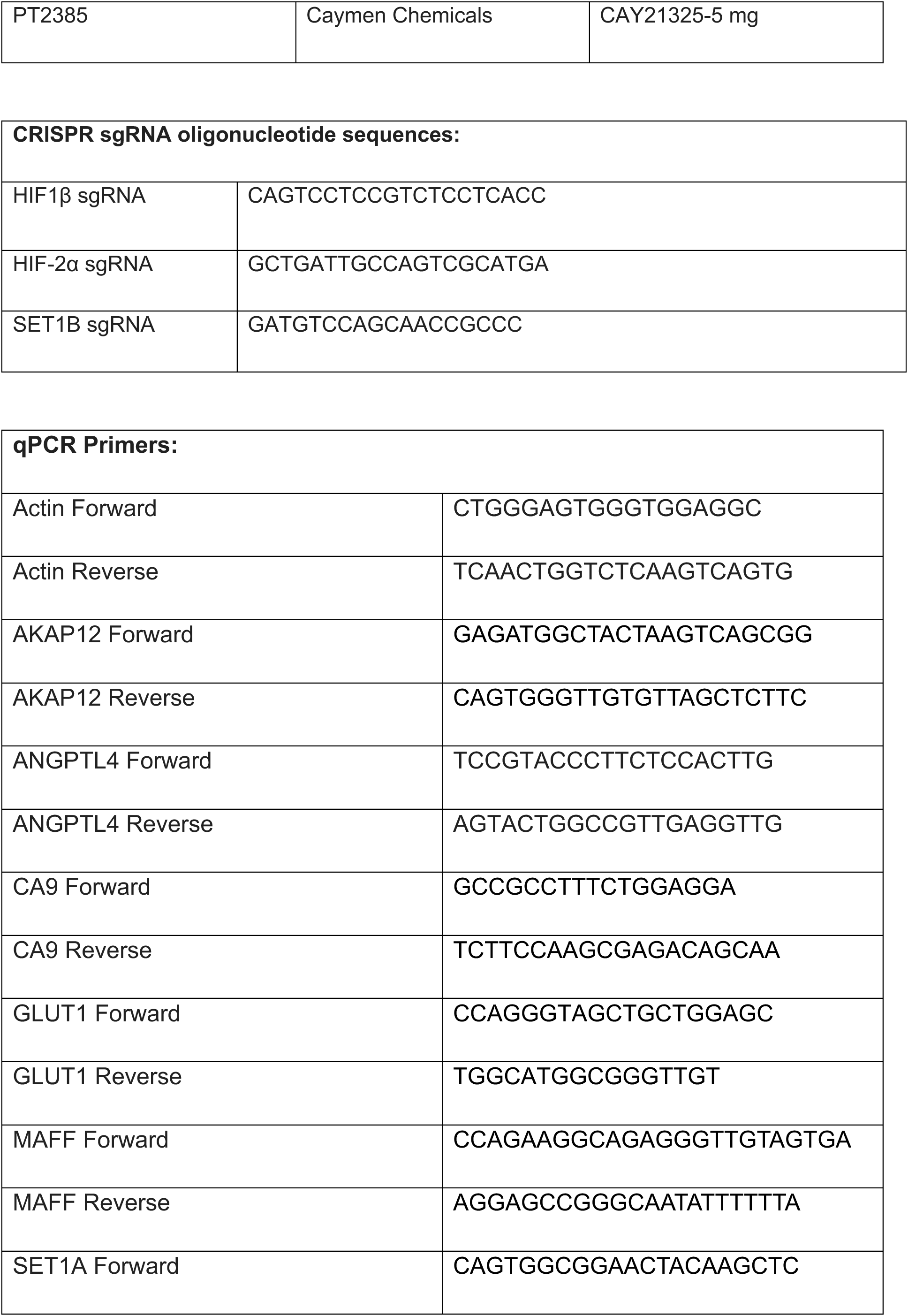
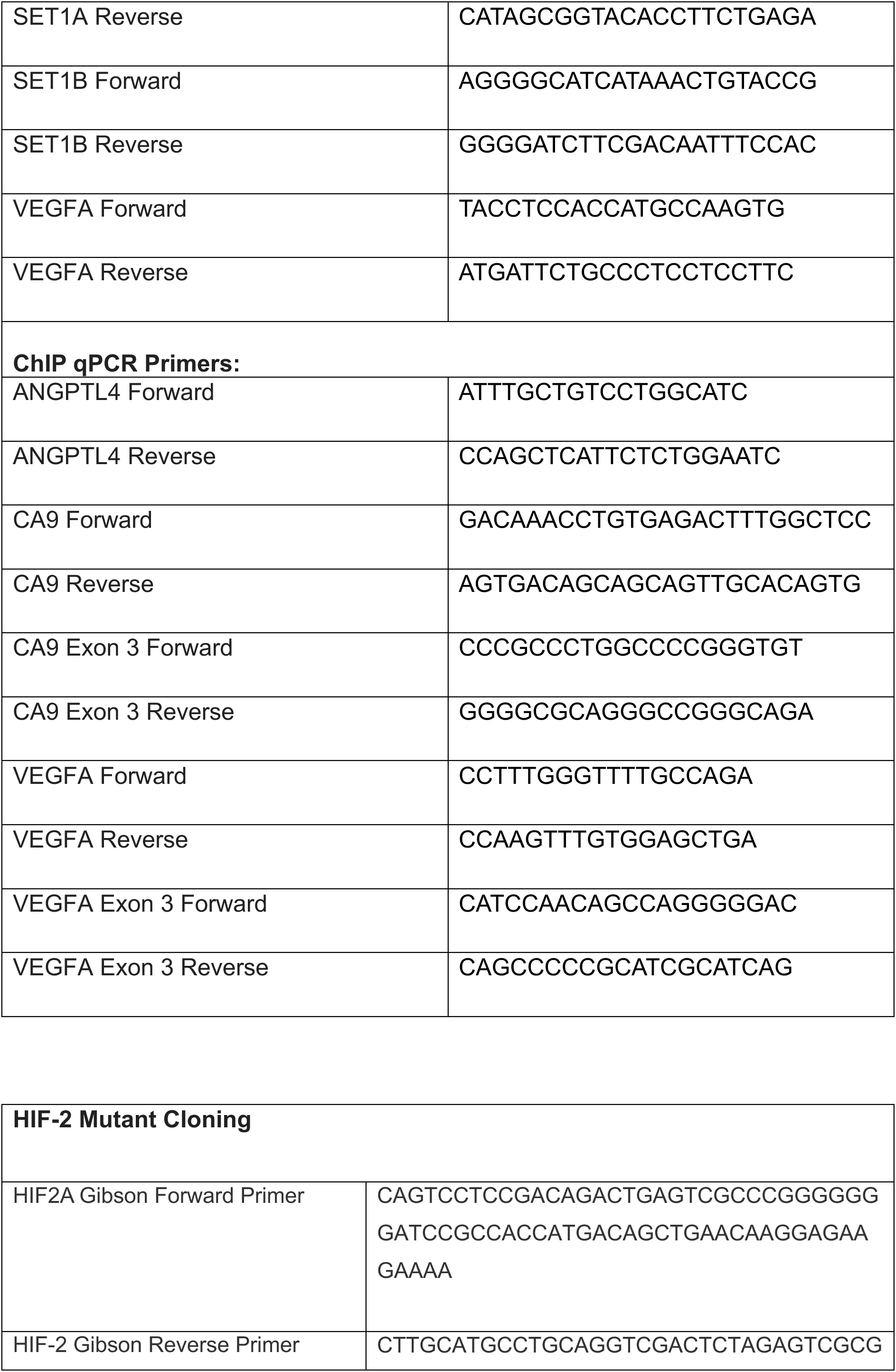
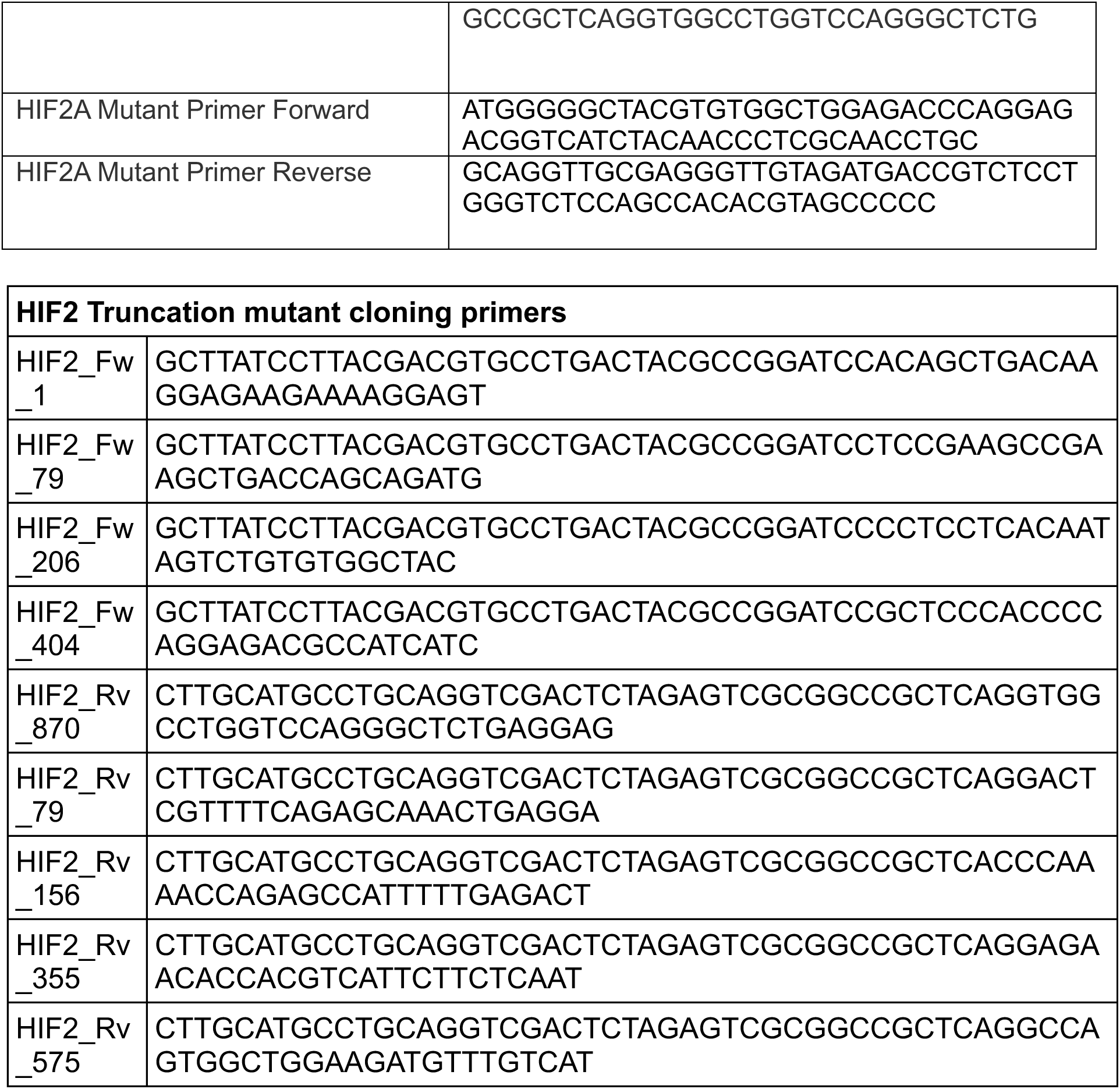

